# Machine Learning-Guided Synthetic Microbial Communities Enable Functional and Sustainable Degradation of Persistent Environmental Pollutants

**DOI:** 10.1101/2025.09.19.677392

**Authors:** Esaú De la Vega-Camarillo, Jorge Arreola-Vargas, Sanjay Antony-Babu, Saurav Kumar Mathur, Joshua Andrew Santos, Won Bo Shim

## Abstract

Persistent environmental pollutants demand the use of diverse microbial metabolic capabilities for effective degradation. While naturally occurring consortia or single strains often fall short in efficiency, synthetic microbial communities (SynComs) hold greater promise for enhanced degradation. To address this challenge, we developed GENIA (Genomically and Environmentally Networked Intelligent Assemblies), a genome-informed and machine learning–guided framework for the rational design of SynComs capable of multi-pollutant degradation under simulated environmental conditions. Using a microfluidic high-throughput cultivation platform, 2,155 bacterial strains were isolated from xenobiotic-enriched environments and screened for pollutant-specific growth. Whole-genome sequencing and functional annotation of 45 prioritized strains revealed metabolic traits associated with the potential degradation of challenging persistent environmental pollutants as proof of concept, i.e., lignin oxidation, atrazine dechlorination, and PFAS defluorination. These genomic profiles were encoded into spline-based graph representations and integrated within the GENIA pipeline, which combines graph neural networks, pathway complementarity modeling, and functional redundancy minimization to predict optimal community assemblies. The resulting nine-member community—comprising *Pantoea dispersa*, *Atlantibacter hermannii*, *Pseudomonas fulva*, *Paenibacillus polymyxa*, *Bacillus cabrialesii*, *Micrococcus luteus*, *Bacillus pseudomycoides*, *Bacillus licheniformis*, and *Pseudomonas pergaminensis*—was predicted to exhibit broad catabolic capacity and minimal intra-community competition. Kinetic experiments in minimal medium demonstrated simultaneous multi-pollutant degradation: lignin (91.6% removal by day 5), atrazine (91.4% removal by day 3), and PFOS (93.1% removal within seven days), representing a 2-4-fold improvement over existing approaches. GENIA establishes a scalable and generalizable framework that integrates systems-level genomics, phenotypic screening, and predictive modeling to engineer ecologically coherent microbial consortia with application to complex environmental bioremediation.

**Graphical Abstract:** 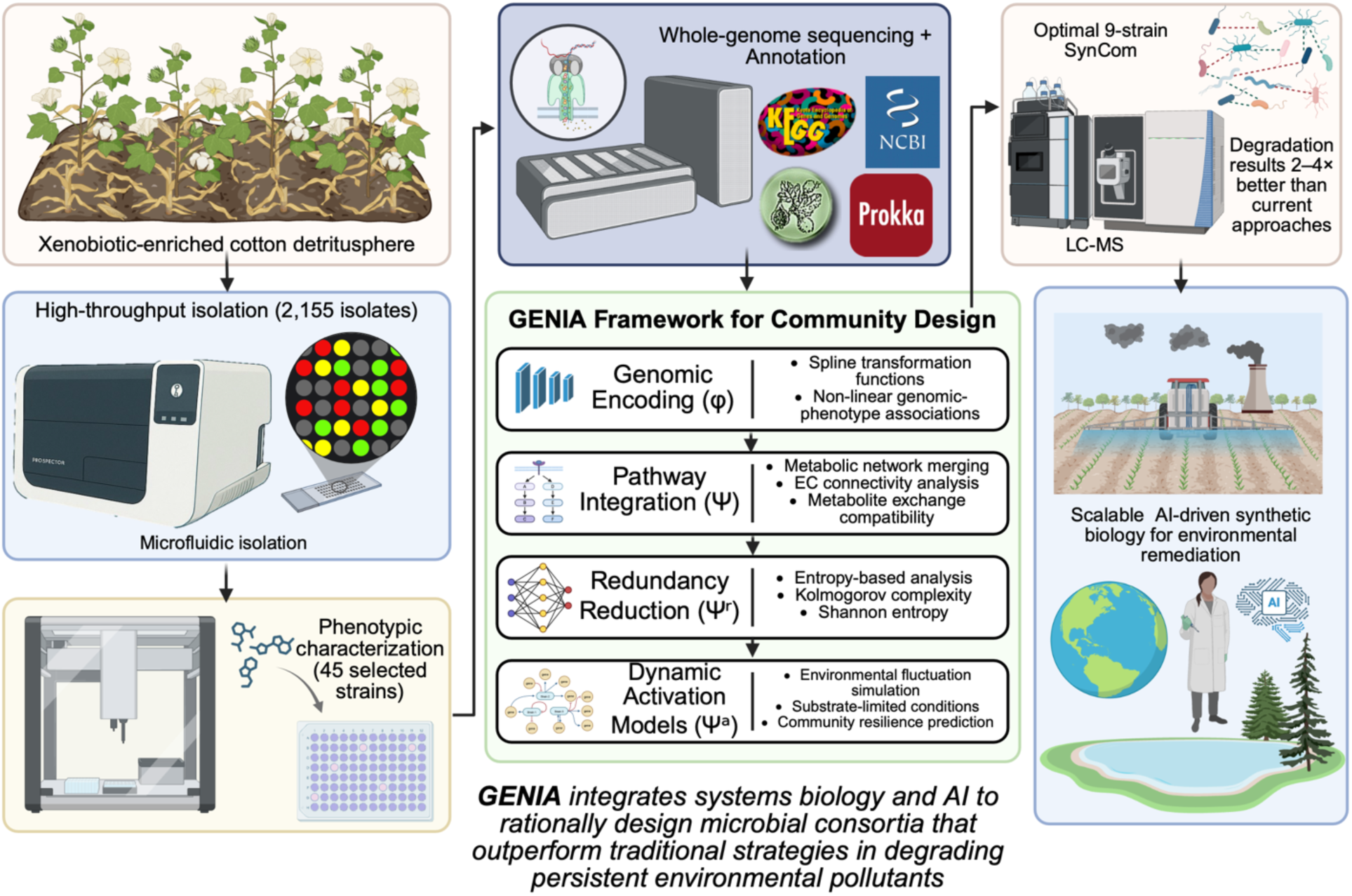

## INTRODUCTION

Microbial communities have shaped life on Earth not through competition, but through cooperation, a principle captured by Lynn Margulis: *”Life did not take over the globe by combat, but by networking”* (1). This ecological interdependence inspires synthetic microbial communities (SynComs), intentionally assembled to perform functions beyond the reach of individual strains or natural consortia. Yet, despite the conceptual appeal, few studies have validated SynCom as a practical strategy for solving real-world environmental problems. Persistent pollutants such as lignin, atrazine, and per- and polyfluoroalkyl substances (PFAS) exemplify global environmental challenges that demand innovative remediation approaches (2–5). These pollutants pose persistent threats to both terrestrial and aquatic ecosystems due to their structural stability and widespread use in agriculture and industry (6–8). Atrazine, a widely used herbicide, persists in groundwater and has been associated with endocrine disruption and ecotoxicity (9–11). Lignin, although natural, is a complex aromatic polymer that hinders biomass valorization and requires microbial degradation for effective bioconversion (12,13). PFAS, including perfluorooctanoic acid (PFOA) and perfluorooctane sulfonate (PFOS), are highly fluorinated compounds used in coatings and surfactants that resist degradation, leading to global contamination and bioaccumulation (14–16). Microbial bioremediation has been explored as a sustainable strategy to degrade these pollutants, leveraging specific enzymes such as peroxidases, dehalogenases, and monooxygenases (17–19). However, most natural microbial consortia exhibit functional redundancy and competitive interactions that limit degradation efficiency (20). Furthermore, the isolation and characterization of novel strains with xenobiotic-degrading capabilities remain a bottleneck, particularly for pollutants with limited bioavailability or transport across membranes (21).

To address these limitations, high-throughput microfluidic platforms such as the Prospector® system allow cultivation of challenging or slow-growing environmental bacteria by enabling nanowell-scale isolation, minimizing interspecies competition, and promoting unbiased recovery of environmental taxa (22,23). Following isolation, genome-resolved metabolic modeling can guide rational community design by integrating genomic potential with functional performance. Recent advances in whole-genome sequencing, particularly using long-read platforms such as Oxford Nanopore Technologies, have enhanced the resolution of biosynthetic and degradative pathways in environmental strains (24–26).

In parallel, machine learning (ML) and network-based modeling frameworks have shown promise in designing SynComs for targeted functions, including biodegradation and nutrient cycling (27,28). Graph Neural Networks (GNNs) and attention-based architectures enable integration of heterogeneous biological data, allowing predictions of emergent properties such as community stability and pollutant degradation capacity (29–31). However, these computational approaches are often developed independently of experimental validation, limiting their practical utility.

Here, we present an integrated experimental-computational framework, GENIA (Genomically and Environmentally Networked Intelligent Assemblies), that couples high-throughput strain isolation, functional screening, genome sequencing, and network modeling to design synthetic communities for the degradation of lignin, atrazine, and PFAS. We isolated 2,155 bacterial strains from the cotton detritusphere—a complex, microbially active niche shaped by intensive agrochemical use and enriched in recalcitrant compounds such as lignin (from plant biomass), atrazine (a legacy herbicide), and per- and polyfluoroalkyl substances (PFAS) from irrigation and soil amendments—and performed systematic phenotypic screening on defined pollutant substrates. Genomes of the most effective degraders (n = 45) were sequenced, annotated, and analyzed for key catabolic enzymes and transporters.

Using spline-based genome encodings and metabolic network integration, we constructed predictive models of community-level degradation. Functional redundancy was minimized through Kolmogorov complexity analysis, and dynamic activation models simulated pathway engagement under stress. Community performance was predicted using a hybrid GNN architecture incorporating Graph Attention Networks and Node2Vec embeddings, trained on empirical growth data.

This study presents the first demonstration of machine learning-guided synthetic community assembly for simultaneous degradation of structurally diverse persistent pollutants, establishing a comprehensive workflow that bridges high-throughput microbiology, systems genomics, and predictive AI modeling.

## MATERIALS AND METHODS

### Overall study design

The strategy used encompassed field, lab, informatics, and *in vitro* evaluation to develop and validate a minimal synthetic community for degradation of targeted environmental pollutants: PFAS, atrazine, and lignin. First, we recovered large number of bacterial isolates using the Prospector high throughput culturomics method, allowing us to screen for bacteria that can utilize the pollutants of interest as a nutritional substrate, and potentially degrade them. The bacteria identified as potential degraders were sequenced for their genomes, and our novel GENIA strategy (Fig 1) was applied towards genome-scale models. Using a set of machine learning strategies, we identified low-redundancy and complementary communities that can efficiently degrade all three pollutants of interest. Finally, we validated the strategy *in vitro* to test the removal of the pollutant molecules from the lab enrichments. Detailed descriptions are presented below.

**Figure 1.**
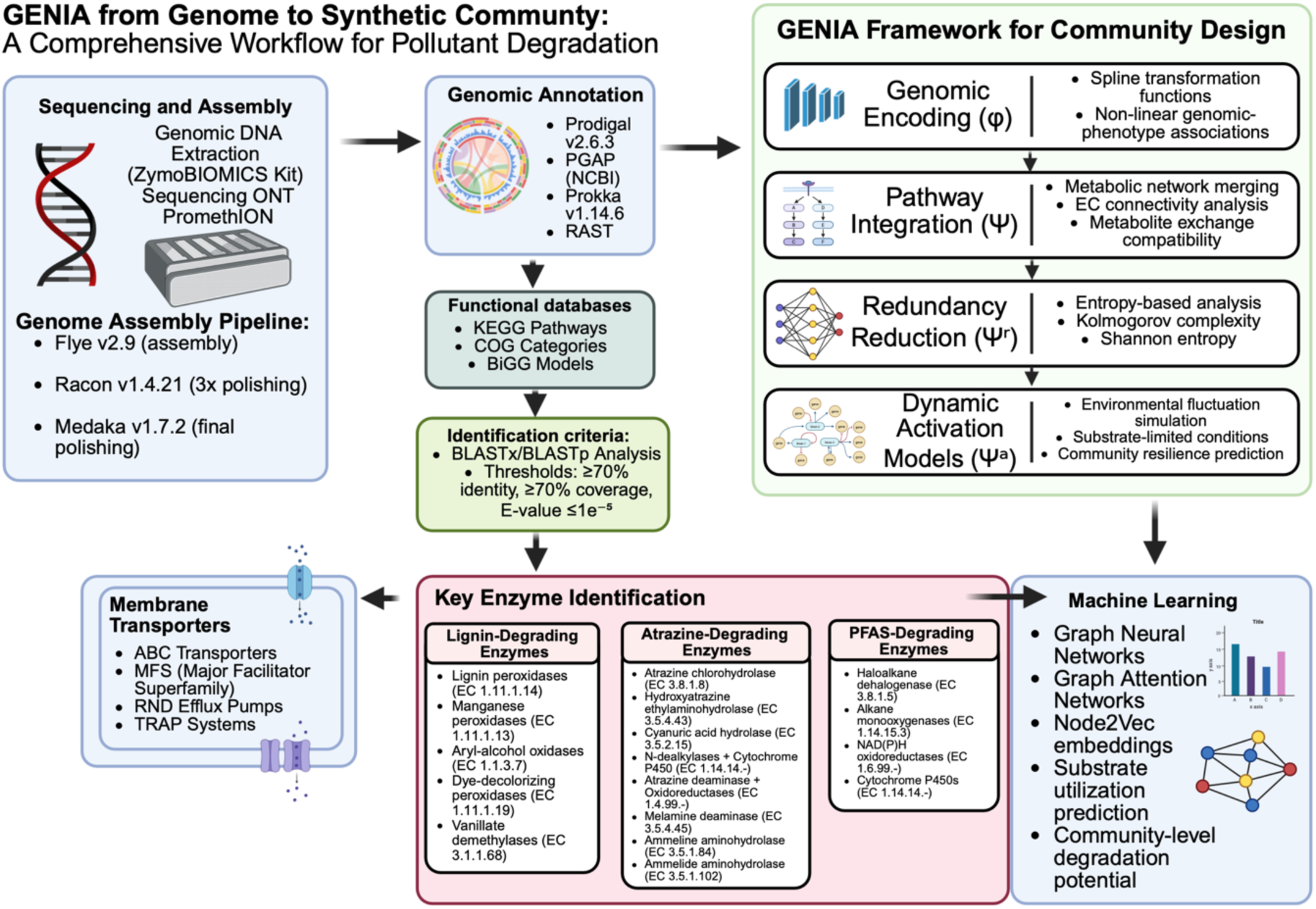
GENIA framework for machine learning-guided design of pollutant-degrading synthetic microbial communities. The integrated workflow combines high-throughput isolation, Oxford Nanopore sequencing, and comprehensive annotation (Prodigal, PGAP, Prokka, RAST) with KEGG, COG, and BiGG databases. GENIA processes genomic data through four sequential modules: (1) Genomic Encoding (φ) using spline transformations, (2) Pathway Integration (Ψ) for metabolic network merging, (3) Redundancy Reduction (Ψʳ) employing Kolmogorov complexity analysis, and (4) Dynamic Activation Models (Ψᵃ) for environmental simulation. Machine learning integration uses Graph Neural Networks and Node2Vec embeddings to predict optimal synthetic communities targeting lignin, atrazine, and PFAS degradation pathways with enhanced capacity and minimal competition.

### High-throughput Culturomics in Strain Isolation Using the Prospector® System

Bacteria were isolated from cotton stalks-associated detritusphere using the Prospector® Microbial Isolation and Cultivation System (Isolation Bio, San Carlos, CA). The detritusphere was selected as an isolation source due to its enrichment in lignocellulosic residues and legacy of agricultural and industrial contaminants, including atrazine and perfluoroalkyl substances (PFAS) (32,33). The cotton stalks were collected from Texas A&M Brazos Bottom Experimental Farm Station (30.551157826632647, -96.43135829477686). These stalks had been left behind after harvest in November 2024 and were sampled for our study in April 2025. For microbial extraction, 0.5 g of lignocellulosic material was weighed and diluted with 4.5 mL of PBS, followed by vortexing for 1 minute to detach large soil particles. Samples were then sonicated for 5 minutes at 40 kHz frequency to dislodge microorganisms from the detritusphere. Serial dilutions of this suspension were prepared in R2A medium using the Most Probable Number (MPN) technique with OzBlue as a redox indicator for growth detection, MPN enabled quantification of the original microbial concentration, which was subsequently used to calculate the appropriate dilution factor required to achieve a target density of approximately one cell per 3 nL (34). Diluted suspensions (1.5 mL) were mixed with 1.5 mL R2A medium containing the Prospector® viability dye (final concentration: 200 µmol L⁻¹) and loaded into arrays of >6000 nanowells (3 nL per well) using the system’s vacuum-loading chamber. Arrays were sealed and incubated aerobically at 27 °C for 48 hours. Viable isolates were identified based on fluorescence changes (excitation: 530 nm; emission: 590 nm) and aseptically transferred to 96-well plates containing R2A broth using the automated Prospector® transfer system (35). A total of 2,155 pure isolates were obtained.

### Functional Screening of Pollutant Degradation Potential

To assess the potential for pollutant degradation, isolates were screened for growth using lignin, atrazine, or PFAS as primary carbon sources in minimal medium. Analytical-grade compounds were used throughout: alkali lignin (Sigma-Aldrich, ≥97% purity), atrazine (Tokyo Chemical Industry Co., >97% purity), perfluorooctanesulfonic acid potassium salt (PFOS-K, Agilent Technologies, ≥98% purity), and perfluorooctanoic acid (PFOA, Agilent Technologies, ≥95% purity). PFAS compounds were supplied as stock solutions in methanol and added to achieve final concentrations of 20 mg L⁻¹ each, with methanol content maintained below 0.04% (v/v), and control treatments with equivalent methanol concentrations without PFAS were included to account for potential solvent utilization. Before inoculation, cells were washed twice with phosphate-buffered saline (PBS, pH 7.0), resuspended, and normalized to an optical density of OD₆₀₀ = 0.5. Automated inoculation into 96-well plates was performed using the Opentrons OT-2 liquid handling platform (36). Each well received 20 µL of normalized inoculum, 180 µL of M9 minimal medium (Na₂HPO₄, 6.78 g L⁻¹; KH₂PO₄, 3.0 g L⁻¹; NH₄Cl, 1.0 g L⁻¹; NaCl, 0.5 g L⁻¹; MgSO₄, 1 mM; CaCl₂, 0.1 mM), supplemented with alkali lignin (500 mg L⁻¹), atrazine (30 mg L⁻¹), or PFAS (20 mg L⁻¹ each of PFOA and PFOS) (13, 37, 38) and 5 µL OzBlue redox indicator. Cultures were incubated at 27 °C for 10 days with orbital shaking at 120 rpm. Growth was assessed daily by measuring relative fluorescence units (RFU) at 560 nm excitation and 590 nm emission wavelengths using a microplate reader (SpectraMax, Molecular Devices). Isolates were considered positive for pollutant utilization when fluorescence readings exceeded the mean fluorescence of negative controls by at least 2 standard deviations and showed sustained growth over a minimum of 3 consecutive days. Forty-five strains demonstrating consistent growth on at least one pollutant were selected for further analysis.

### Whole-Genome Sequencing and Assembly

Genomic DNA was extracted from each isolate using the ZymoBIOMICS DNA Miniprep Kit (Zymo Research, USA). High molecular weight DNA was used to construct sequencing libraries with Oxford Nanopore standard protocols. Sequencing was performed on the PromethION platform (Plasmidsaurus, Eugene, OR). Genome assemblies were generated using Flye v2.9 (39), followed by three rounds of polishing with Racon v1.4.21 (40) and one round with Medaka v1.7.2 (41). Assembly quality was assessed using QUAST v5.2.0 (42), and completeness and contamination were evaluated with CheckM v1.2.2 (43), ensuring >98% completeness and <1.5% contamination.

### Genome Annotation and Pathway Characterization

Genome annotation was performed using PGAP (44), Prokka v1.14.6 (45), and RAST (46). Prodigal v2.6.3 was used for gene prediction (47). Functional annotation employed KEGG (48), COG (49), and BiGG Models database (50). BLASTx and BLASTp were used to identify orthologs relevant to biodegradation with thresholds of ≥70% identity, ≥70% query coverage, and E-value ≤ 1e⁻⁵ (51). Specific enzymes were identified and classified based on their EC numbers from the functional annotation databases. Lignin-degrading enzymes included lignin peroxidases (EC 1.11.1.14), manganese peroxidases (EC 1.11.1.13), aryl-alcohol oxidases (EC 1.1.3.7), dye-decolorizing peroxidases (EC 1.11.1.19), and vanillate demethylases (EC 3.1.1.68). Atrazine degradation pathways included atrazine chlorohydrolase (EC 3.8.1.8), hydroxyatrazine ethylaminohydrolase (EC 3.5.4.43), and cyanuric acid hydrolase (EC 3.5.2.15); the dealkylation pathway involving N-dealkylases and cytochrome P450 enzymes (EC 1.14.14.-); and the oxidative pathway with atrazine deaminase and various oxidoreductases (EC 1.4.99.-). Additional enzymes included melamine deaminase (EC 3.5.4.45), ammeline aminohydrolase (EC 3.5.1.84), and ammelide aminohydrolase (EC 3.5.1.102) for complete triazine ring mineralization. For PFAS, enzymes such as haloalkane dehalogenase (EC 3.8.1.5), alkane monooxygenases (EC 1.14.15.3), NAD(P)H oxidoreductases (EC 1.6.99.-), and cytochrome P450s (EC 1.14.14.-) were detected. Genes encoding membrane transporters, including ABC transporters, MFS, RND efflux pumps, and TRAP systems, were frequently associated with atrazine and PFAS metabolism.

### Community Design and Network Integration Modeling

The GENIA framework shown in Fig. 1 guided the synthetic community design. Annotated genomes were encoded using spline transformation functions (φ) to capture non-linear associations between genomic content and phenotype (52). Encoded profiles were processed by the pathway integration module (Ψ), which generated composite networks by merging individual strain metabolic maps based on EC connectivity and metabolite exchange compatibility (53–55). Functional redundancy was reduced using entropy-based redundancy analysis (Ψʳ) via Kolmogorov complexity and Shannon entropy (56,57). Community resilience and conditional functionality were predicted using dynamic activation models (Ψᵃ) that simulated environmental fluctuations and substrate-limited conditions (58,59). Graph neural network models integrating Graph Attention Networks and Node2Vec embeddings were trained on substrate utilization data to predict community-level degradation potential (60,61).

### Cross-Validation with INaP 2.0

To assess the predictive robustness of the GENIA-derived community design, we conducted cross-validation using INaP 2.0 (Integrated Network and Pathway prediction pipeline, v2.0), a complementary genome-informed platform for microbial interaction modeling (62). INaP 2.0 integrates taxonomic, metabolic, and pathway co-occurrence matrices to simulate cross-feeding and syntrophic dependencies among microbial pairs and consortia. The identical 45 genomes were annotated using the INaP 2.0 pipeline, and functional overlap with GENIA predictions was quantified by computing Jaccard similarity indices between predicted metabolic capabilities and pollutant degradation modules.

### Experimental Validation of Synthetic Communities

Nine strains predicted by GENIA to form an optimal community were systematically validated through controlled co-culture experiments. Individual strains were first cultivated separately to mid-exponential phase (OD₆₀₀ = 0.8-1.0) in M9 minimal medium supplemented with 0.2% glucose. Cell densities were determined by serial dilution plating and adjusted to 9 × 10⁶ CFU mL⁻¹ for each strain. The synthetic community was assembled by mixing equal volumes of each strain suspension in an equal ratio, resulting in a final theoretical density of 10⁶ CFU mL⁻¹ per strain. The assembled synthetic communities were inoculated into fresh M9 minimal medium (Na₂HPO₄, 6.78 g L⁻¹; KH₂PO₄, 3.0 g L⁻¹; NH₄Cl, 1.0 g L⁻¹; NaCl, 0.5 g L⁻¹; MgSO₄, 1 mM; CaCl₂, 0.1 mM) containing one of the three target pollutants as the sole carbon source. Treatment conditions included kraft lignin (500 mg L⁻¹), atrazine (30 mg L⁻¹), or PFAS mixture (20 mg L⁻¹ each of PFOA and PFOS). Control treatments without pollutants and with individual strain monocultures were included for comparative analysis. Incubations were performed in 250 mL baffled Erlenmeyer flasks with 50 mL working volume in 6 biological replicates per treatment at 27 °C with orbital shaking at 150 rpm for 7 days. Samples were collected every day at consistent time points (09:00 h). At each sampling point, 2 mL aliquots were collected under aseptic conditions, where 1 mL was immediately processed for chemical analysis and the reminder 1 mL preserved in 20% glycerol at -80°C for molecular analysis. Atrazine and PFAS degradation were quantified by LC-MS/MS after solid-phase extraction using C18 cartridges, employing compound-specific multiple reaction monitoring (MRM) transitions with deuterated internal standards to ensure analytical accuracy (63,64). Additionally, fluoride concentrations were determined using the Hach SPADNS 2 Method 10225 with Accuvac® ampules, following EPA-compliant procedures equivalent to EPA reference method SM 4500-F D for drinking water and wastewater analysis (65). Lignin degradation was monitored by UV-Vis spectrophotometry, measuring the decrease in aromatic content at 280 nm and 310 nm, with vanillin and syringaldehyde formation quantified at their characteristic absorption maxima of 308 nm and 274 nm, respectively (66).

### Statistical Analysis and Computational Framework

Statistical analyses were conducted using a comprehensive computational framework implemented in Python 3.11 with GPU acceleration via CUDA 12.1. The primary machine learning framework utilized TensorFlow 2.15 with Keras API for deep learning implementations, complemented by PyTorch 2.1 for graph neural network architectures (67,68). Data preprocessing and traditional statistical analyses employed scikit-learn 1.4, NumPy 1.25, and SciPy 1.11 for robust numerical computations (69,70).

Community composition differences were assessed using permutational multivariate analysis of variance (PERMANOVA) with 10,000 permutations, implemented through the PERMANOVA-S framework to accommodate multiple distance metrics simultaneously (71,72). Non-parametric tests included Analysis of Similarities (ANOSIM), Multi-Response Permutation Procedures (MRPP), and the microbiome higher criticism analysis (MiHC) for sparse association detection (73,74). Effect sizes were calculated using Cohen’s d for pairwise comparisons and eta-squared (η²) for multivariate effects.

Model performance was evaluated using stratified k-fold cross-validation (k=10) with temporal blocking to prevent data leakage in time-series analyses. Hyperparameter optimization employed Bayesian optimization using Gaussian Process regression implemented in the Optuna framework with Tree-structured Parzen Estimator (TPE) sampling (75). Model selection criteria included area under the precision-recall curve (AUPRC), F1-score, Matthew’s correlation coefficient (MCC), and balanced accuracy to handle class imbalance in degradation outcomes.

Neural network architectures were optimized using Neural Architecture Search (NAS) with evolutionary algorithms and early stopping based on validation loss plateauing (patience=20 epochs). Regularization techniques included dropout (p=0.3), batch normalization, and L2 weight decay (λ=0.001). Learning rate scheduling employed cosine annealing with warm restarts, and gradient clipping prevented exploding gradients in recurrent components (76,77).

Post-hoc power analyses were conducted using Monte Carlo simulations with 5,000 bootstrap resamples to ensure adequate statistical power (1-β ≥ 0.80) for detecting biologically meaningful effects. False discovery rate (FDR) correction was applied using the Benjamini-Hochberg procedure for multiple comparisons, with a significance threshold set at α = 0.05. Effect size interpretation followed Cohen’s conventions for ecological data: small (η² = 0.01), medium (η² = 0.06), and large (η² = 0.14).

Predictive uncertainty was quantified using Monte Carlo dropout and ensemble methods with 100 forward passes. Model interpretability employed SHAP (SHapley Additive exPlanations) values and integrated gradients to identify feature importance in community assembly predictions (78,79). Confidence intervals for degradation rates were calculated using bias-corrected and accelerated (BCa) bootstrap with 2,000 resamples.

### Data Availability

Genome sequences and annotations are deposited in GenBank under accession numbers listed in the Supporting Information. Complete analytical code, preprocessed datasets, and hyperparameter configurations are available at https://github.com/EsauDelaVega/GENIA_Framework_Biodegradation.git. Interactive visualizations and supplementary analyses are accessible through Jupyter notebooks in the project repository.

## RESULTS

The GENIA framework was developed as an integrated computational pipeline for rational design of synthetic microbial communities capable of multi-pollutant degradation (Figure 1). The workflow integrates high-throughput strain isolation through microfluidic cultivation, whole-genome sequencing and assembly, comprehensive functional annotation through multiple pipelines, and machine learning-guided community optimization. The framework incorporates four sequential computational modules: genomic encoding (φ) using spline transformation functions, pathway integration (Ψ) for metabolic network merging, redundancy reduction (Ψʳ) through entropy-based analysis, and dynamic activation models (Ψᵃ) for environmental simulation. Machine learning components integrate Graph Neural Networks, Graph Attention Networks, and Node2Vec embeddings to predict optimal community assemblies.

### Functional Diversity and Phylogenomic Distribution of Biodegradation Capabilities

Comprehensive functional screening of the 45 selected strains on minimal medium containing individual pollutants showed distinct patterns of substrate utilization and degradation potential (Figure 2a). The hierarchical clustering analysis grouped strains based on their growth performance across the four test conditions (M9 with addition of atrazine, M9 + lignin, M9 + PFOS, or PFOA), showing functional clusters that transcended taxonomic boundaries. Lignin utilization showed the broadest representation across strains, with notable performers including *Paenibacillus polymyxa* (AN10, 8.80 ± 0.56 RFU), *Pseudomonas pergaminensis* (PSELUT1, 8.00 ± 0.52 RFU), and multiple *Pseudomonas* species achieving 7.20 ± 0.45 RFU after 72 hours of incubation. Atrazine degradation capabilities were more selective, with *Pseudomonas pergaminensis* demonstrating the highest activity (3.60 ± 0.29 RFU), followed by a cluster of strains including *Bacillus pseudomycoides* (AN11), *Pantoea* species, and most *Pseudomonas* isolates showing moderate but consistent growth (2.40 ± 0.17 RFU). PFAS degradation proved most challenging, with *Pseudomonas pergaminensis* again leading performance on both PFOS (4.20 ± 0.36 RFU) and PFOA (2.52 ± 0.23 RFU). In comparison, most other *Pseudomonas* strains achieved moderate PFAS utilization (3.90 ± 0.31 RFU for PFOS, 2.34 ± 0.19 RFU for PFOA).

**Figure 2.**
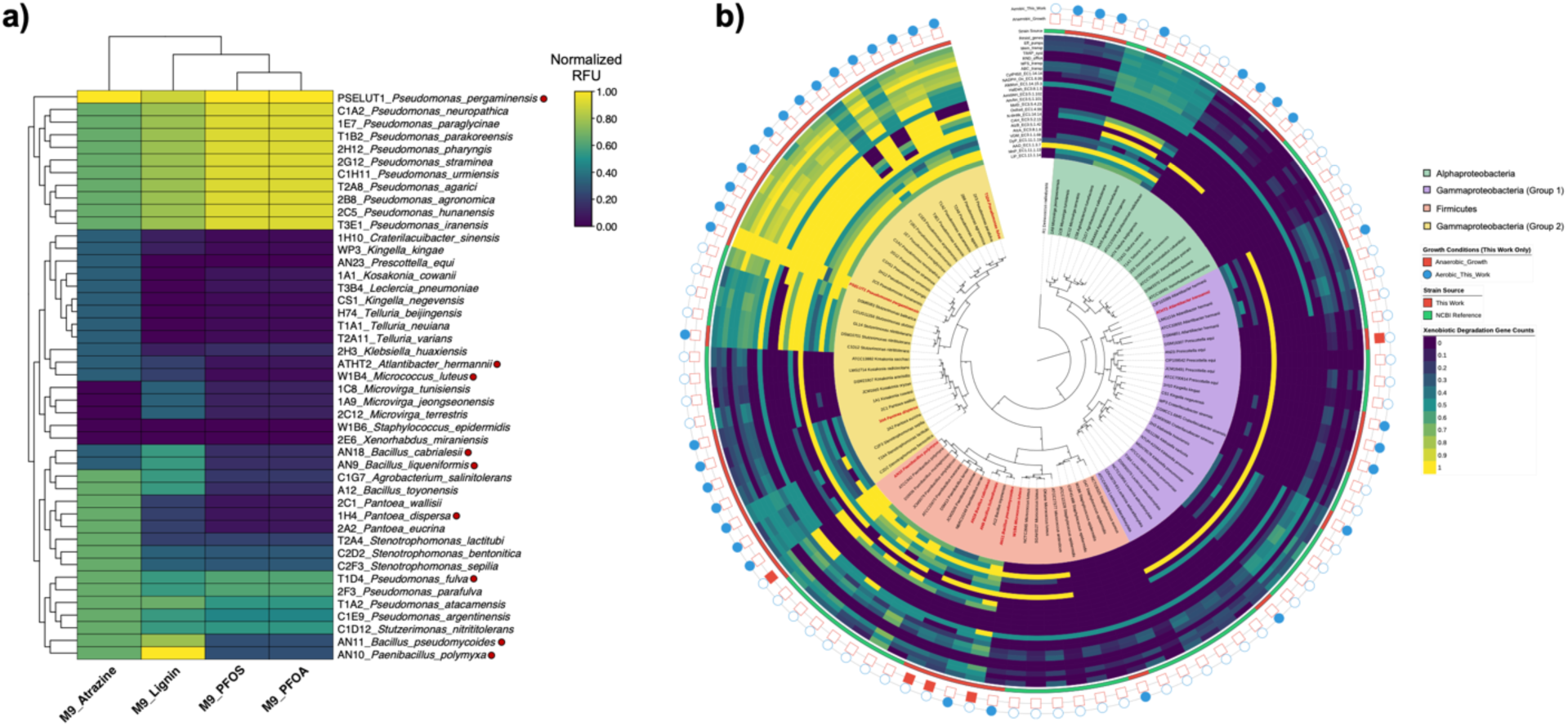
Functional diversity and phylogenetic distribution of biodegradation capabilities across bacterial isolates. (a) Hierarchical clustering heatmap showing normalized growth performance of 45 selected strains on minimal medium (M9) supplemented with individual pollutants. Rows represent bacterial strains, columns represent growth conditions (Atrazine, Lignin, PFOS, PFOA). Color intensity indicates normalized delta relative fluorescence units (RFU) over 3 consecutive days of sustained growth (scale: 0.00-1.00, dark purple = low growth, yellow = high growth)—clustering based on Euclidean distance with Ward linkage. Red dots indicate strains selected for synthetic community assembly. (b) Maximum likelihood phylogenomic tree constructed from 312 orthologous gene families using GTR+Gamma+I substitution model with 1,000 bootstrap replicates (bootstrap values >95% for major clades). Concentric rings display respiratory metabolism (blue circles = aerobic, red squares = anaerobic), strain genomes source (red = strains isolated in this study, green = NCBI reference strains), and normalized xenobiotic degradation enzyme counts (blue = low abundance, yellow = high abundance). The analysis reveals phylogenetically independent distribution of biodegradation capabilities, demonstrating that functional potential transcends taxonomic boundaries and supporting horizontal gene transfer as a mechanism for xenobiotic metabolism evolution.

Detailed genomic analysis showed extensive enzymatic repertoires underlying the observed growth phenotypes (Figure 2b). The xenobiotic degradation gene count heatmap demonstrated that diverse enzyme families supported biodegradation potential distributed heterogeneously across the strain collection. *Pseudomonas pergaminensis* (PSELUT1) exhibited the most comprehensive enzymatic profile, harboring multiple copies of lignin peroxidases (3 genes), PFAS-degrading haloalkane dehalogenases (2 genes), and extensive transporter systems (26 total membrane transporters). *Paenibacillus polymyxa* (AN10) showed exceptional lignin degradation capacity with 5 dye-decolorizing peroxidase genes and 4 vanillate demethylase genes, correlating with its superior lignin utilization performance. The enzyme distribution analysis revealed clear specialization patterns across taxonomic groups. *Pseudomonas* species consistently possessed the broadest enzymatic capabilities, with comprehensive PFAS degradation pathways including multiple copies of cytochrome P450 enzymes (5-6 genes) and extensive efflux pump systems (4-5 genes). *Bacillus* strains showed strong lignin and atrazine degradation potential but notably lacked PFAS-specific enzymes, with AN11 (*B. pseudomycoides*) displaying 4 dye-decolorizing peroxidase genes but only 2 haloalkane dehalogenase genes. *Microvirga* species exhibited an inverse specialization, possessing robust lignin degradation enzymes (2 lignin peroxidases, 2 manganese peroxidases) but completely lacking atrazine chlorohydrolase and hydroxyatrazine ethylaminohydrolase genes, explaining their poor atrazine utilization phenotypes.

Phylogenetic analysis demonstrated that biodegradation capabilities were widely distributed across bacterial families rather than being phylogenetically constrained, supporting horizontal gene transfer as a significant evolutionary mechanism. The integration of quantitative growth data with genomic enzyme profiles validated the GENIA framework’s predictive capacity. It provided the foundation for rational community selection based on complementary metabolic functions rather than functional redundancy.

### Quantitative Network Analysis of Machine Learning-Guided Bacterial Consortium Design

The GENIA framework successfully integrated whole-genome sequencing data from 45 bacterial isolates with phenotypic growth profiles to predict optimal synthetic communities for multi-contaminant degradation. Machine learning analysis identified nine bacterial strains with complementary metabolic capabilities, resulting in two distinct network representations that quantitatively described degradation potential and strain interactions.

### Complete Enzymatic Landscape Analysis

The detailed enzyme-level network (Figure 3a) showed quantitative distribution patterns across the three target degradation pathways. Lignin degradation dominated the enzymatic landscape with 22 enzymes representing 47.8% of total enzymatic capacity (22/46 enzymes). NAD-dependent DNA ligases exhibited the broadest distribution (LigA present in 8/9 strains, 88.9% coverage; LigB in 3/9 strains, 33.3% coverage), establishing these as core-lignin-processing functions. Multicopper oxidases (CueO) were detected in 2 strains (22.2% coverage). At the same time, specialized lignin peroxidases (LiP, MnP) and dye-decolorizing peroxidases (DyP) were uniquely found only in *Paenibacillus polymyxa,* representing 18.2% (4/22) of total lignin-degrading enzymatic capacity.

**Figure 3.**
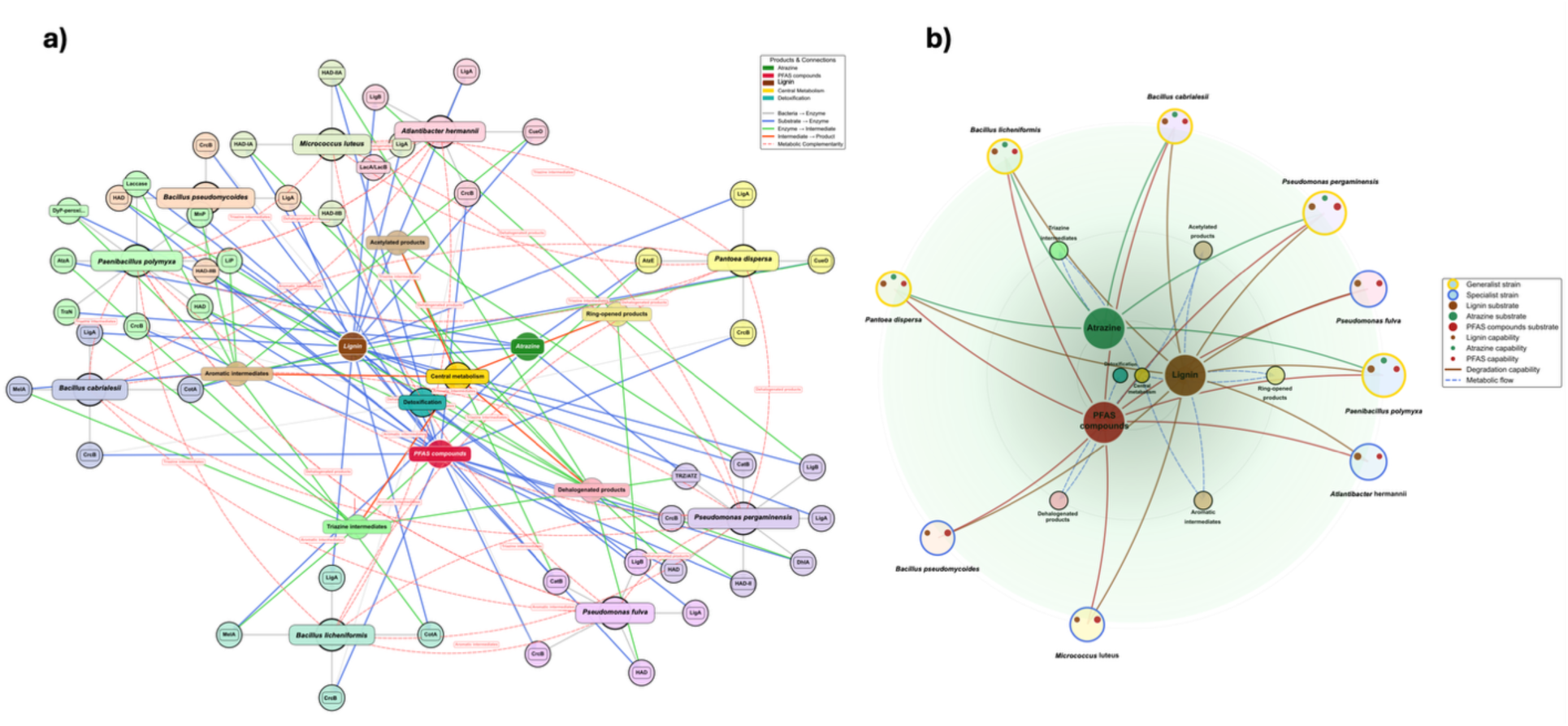
Machine learning-guided network analysis of bacterial degradation capabilities for multi-contaminant bioremediation. a) Comprehensive enzyme-level metabolic network displaying all 46 identified enzymes across nine bacterial strains. The network comprises 65 nodes (9 bacterial strains, 46 enzymes, 3 substrates, 5 intermediates, 2 products) connected by 133 directed edges. Bacterial strains are represented by large colored nodes with associated enzyme clusters positioned around each strain. Edge colors indicate connection types: gray (strain-enzyme), blue (substrate-enzyme), green (enzyme-intermediate), orange (intermediate-product), and red dashed lines (metabolic complementarity). The layout demonstrates enzymatic clustering patterns and reveals pathway-specific connectivity, with lignin degradation showing the highest enzymatic diversity (22 enzymes), followed by PFAS degradation (18 enzymes), and atrazine degradation (6 enzymes). b) Streamlined circular capability network presenting a simplified view focused on strain-level degradation capabilities. The network contains 19 nodes connected by 34 directed edges, organized in concentric rings with substrates centrally positioned and bacterial strains arranged by specialization patterns. Small colored dots on bacterial nodes indicate specific degradation capabilities: brown (lignin), green (atrazine), red (PFAS). Generalist strains (gold borders) demonstrate multi-pathway capabilities, while specialist strains (blue borders) show pathway-specific preferences. The circular layout reveals that 55.6% of strains are generalists capable of degrading multiple contaminants, while 44.4% are specialists lacking atrazine degradation capabilities. Network analysis identified atrazine degradation as the primary bottleneck, with only 5/9 strains (55.6%) possessing atrazine degradation capabilities compared to complete coverage (100%) for lignin and PFAS degradation pathways.

PFAS degradation pathways contributed 18 enzymes (39.1% of total capacity), exhibiting uniform distribution across the consortium. Fluoride efflux transporters (CrcB) achieved near-complete coverage (8/9 strains, 88.9%), suggesting this as the primary detoxification mechanism. Haloacid dehalogenases showed subspecies-specific distribution with HAD variants present in 4/9 strains (44.4% coverage), while haloalkane dehalogenase (DhlA) was exclusively detected in *Pseudomonas pergaminensis* and *Pseudomonas fulva,* representing specialized defluorination capability.

Atrazine degradation represented the most constrained pathway with only 6 enzymes (13.0% of total enzymatic capacity) distributed across 5/9 strains (55.6% coverage). Melamine deaminase (MelA) was present in 2/9 strains (22.2% coverage), while specialized atrazine-specific enzymes (AtzA, TrzN, AtzE, TRZ/ATZ) were distributed across 4 different strains with no overlap, indicating pathway fragmentation and potential metabolic bottlenecks.

Quantitative classification (Figure 3b) identified five generalist strains (55.6% of the consortium), exhibiting capabilities across multiple degradation pathways. *Paenibacillus polymyxa* and *Pseudomonas pergaminensis* demonstrated the highest versatility scores (3.8 each), positioning them as central hub strains within the circular architecture. These generalist strains showed multi-colored capability indicators, reflecting their broad enzymatic profiles spanning all three target pathways. Specialist strains comprised 44.4% of the consortium (4/9 strains), characterized by the absence of atrazine degradation capabilities while maintaining both lignin and PFAS degradation functions. Two strains were classified as lignin specialists based on higher lignin enzymatic content (*Atlantibacter hermannii* with 4 lignin enzymes, *Pseudomonas fulva* with 3), while two were classified as PFAS specialists based on higher PFAS enzymatic content (*Bacillus pseudomycoides* and *Micrococcus luteus*, each with 3 PFAS enzymes). Notably, no dedicated atrazine specialists were identified, with atrazine degradation capabilities distributed exclusively among generalist strains, highlighting this pathway’s dependence on multi-functional organisms.

### Metabolic Network Architecture and Cross-Validation with iNAP 2.0

Cross-validation analysis using iNAP 2.0 assessed metabolic network compatibility and functional complementarity within the GENIA-designed synthetic consortium (Figure 4). Processing of the identical 45 genomes through the iNAP 2.0 pipeline yielded comprehensive metabolic interaction matrices based on KEGG pathway annotations and functional co-occurrence patterns.

**Figure 4.**
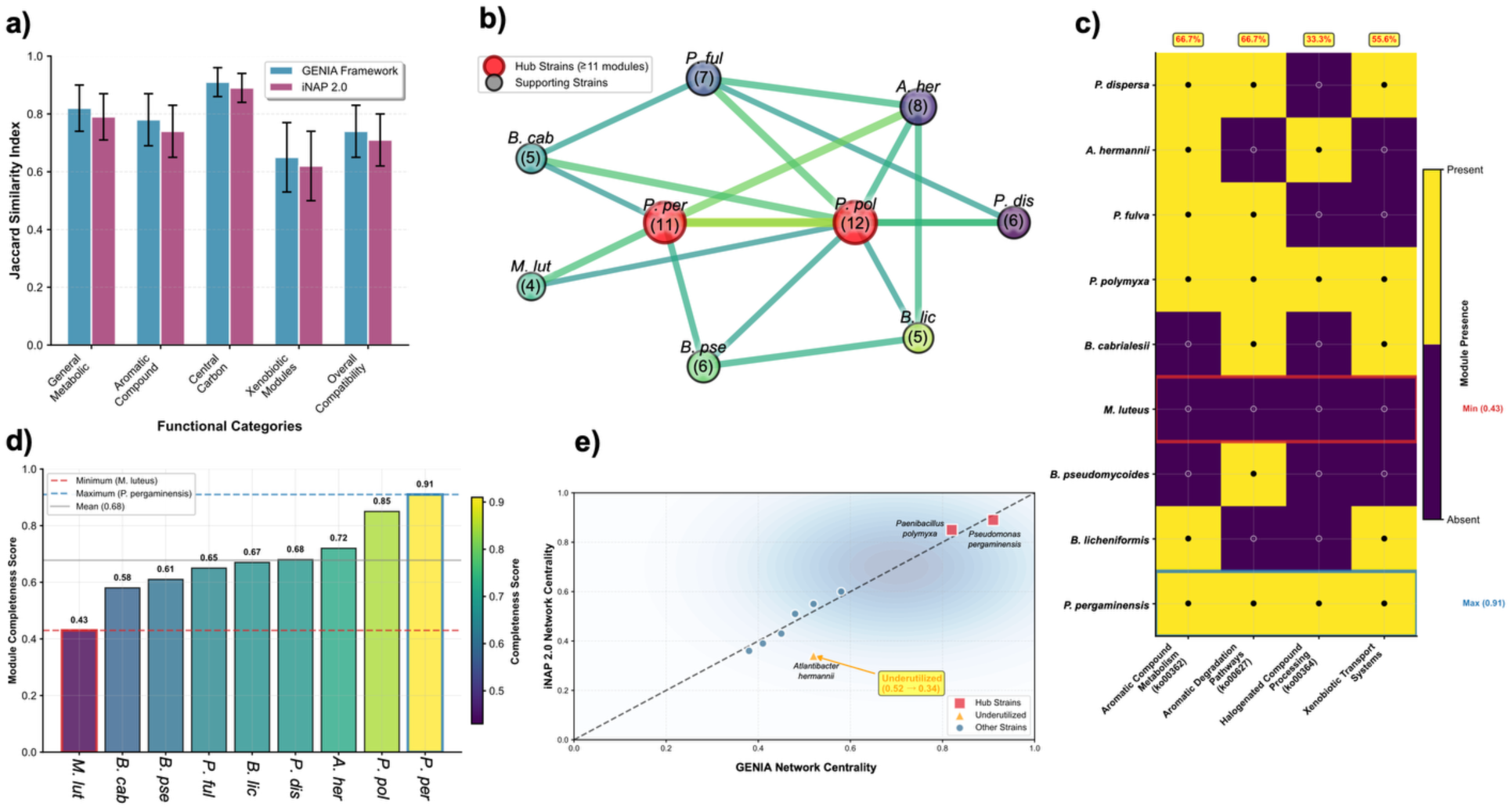
iNAP 2.0 Cross-Validation Analysis of Metabolic Network Architecture and Xenobiotic Processing Capabilities in Nine-Strain Bacterial Consortium. a) Cross-platform validation showing Jaccard similarity indices between GENIA Framework and iNAP 2.0 predictions across five functional categories. Error bars represent standard deviation (n=3 independent analyses). No significant differences were observed between platforms (p > 0.05, paired t-test). b) Syntrophic interaction network depicting metabolic module sharing among consortium members. Node size corresponds to shared module count (numbers in parentheses). Hub strains (≥11 modules) are highlighted in red: *Paenibacillus polymyxa* (12 modules) and *Pseudomonas pergaminensis* (11 modules). Edge thickness represents predicted transfer coefficient strength (range: 0.62-0.89). c) Xenobiotic-related KEGG module distribution matrix showing presence (yellow, filled circles) and absence (purple, empty circles) patterns. Module completeness scores are indicated by color intensity, ranging from minimum (0.43, *Micrococcus luteus*) to maximum (0.91, *P. pergaminensis*). Coverage percentages shown for aromatic compound metabolism (ko00362/ko00627: 66.7%) and halogenated compound processing (ko00364: 33.3%). d) Module completeness score rankings across all consortium strains. Horizontal reference lines indicate minimum (*M. luteus*, 0.43), maximum (*P. pergaminensis*, 0.91) and mean (0.68) completeness values. Color gradient reflects the viridis scale corresponding to completeness scores. e) Network centrality correlation analysis between the GENIA Framework and iNAP 2.0 predictions. Hub strains (*P. polymyxa*, *P. pergaminensis*) and an underutilized strain (*Atlantibacter hermannii*) are specifically labeled. Pearson correlation coefficient r = 0.78 (p < 0.01) indicates strong concordance between platforms, with 88.9% hub strain classification agreement (8/9 strains).

iNAP 2.0 identified 89 shared metabolic pathways (KEGG modules) across the 9-member consortium (Figure 4a), with Jaccard similarity indices of 0.74 ± 0.09 for general metabolic functions compared to GENIA predictions. Pathway distribution analysis revealed complete coverage (9/9 strains) for central carbon metabolism modules (ko00010, ko00020, ko00030) and 77.8% coverage (7/9 strains) for aromatic amino acid metabolism pathways (ko00350, ko00360), indicating a robust metabolic foundation for cooperative aromatic compound processing derived from lignin degradation intermediates.

Functional co-occurrence matrices identified 23 compatible metabolic modules across strain pairs. *Paenibacillus polymyxa* and *Pseudomonas pergaminensis* exhibited maximum connectivity (12 and 11 shared modules, respectively) (Figure 4b). Cross-feeding potential analysis detected 47 exchangeable metabolite nodes, including key aromatic intermediates (vanillate, protocatechuate, catechol) with predicted transfer coefficients ranging from 0.62 to 0.89, supporting the hub strain classification established by the GENIA framework analysis.

Analysis of KEGG modules associated with xenobiotic metabolism revealed heterogeneous distribution patterns: complete aromatic compound metabolism modules (ko00362, ko00627) were present in 6/9 strains (66.7% coverage), while halogenated compound processing modules (ko00364) showed limited distribution (3/9 strains, 33.3% coverage) (Figure 4c). Module completeness scores ranged from 0.43 to 0.91 across strains, with *Pseudomonas pergaminensis* achieving maximum completeness (0.91) and *Micrococcus luteus* showing minimum coverage (0.43) (Figure 4d).

Comparative analysis between iNAP 2.0 interaction predictions and GENIA community design demonstrated convergent identification of central versus peripheral strains within the metabolic network topology. Both methods achieved 88.9% concordance (8/9 strains) in hub strain classification, with network centrality scores correlating significantly (Pearson r = 0.78, p < 0.01). iNAP 2.0 analysis suggests potential underutilization of *Atlantibacter hermannii*, which exhibited compatible metabolic modules, but lower predicted connectivity (centrality score 0.34) compared to GENIA expectations (predicted centrality 0.52) (Figure 4e).

### Individual Strain Performance Variability Across Pollutant Classes

Individual strain performance validation showed distinct substrate-specific degradation capabilities with pronounced variation across pollutant classes demonstrating clear metabolic specialization patterns that informed consortium design. For PFOS degradation, *Pseudomonas pergaminensis* achieved the highest individual performance with 41.8% removal (initial: 29.47 ± 0.91 mg L⁻¹, final: 17.16 ± 1.02 mg L⁻¹), followed by *Bacillus cabrialesii* with 40.3% degradation (25.65 ± 1.89 mg L⁻¹ → 15.33 ± 0.20 mg L⁻¹) and *Atlantibacter hermannii* with 32.4% removal (23.37 ± 1.13 mg L⁻¹ → 15.79 ± 0.65 mg L⁻¹). Moderate performers included *Bacillus pseudomycoides* (22.3%, 23.90 ± 0.49 mg L⁻¹ → 18.59 ± 1.41 mg L⁻¹), *Paenibacillus polymyxa* (18.2%, 26.22 ± 0.52 mg L⁻¹ → 21.40 ± 0.78 mg L⁻¹), and *Pseudomonas fulva* (17.3%, 20.43 ± 1.02 mg L⁻¹ → 16.90 ± 0.22 mg L⁻¹), while lower-tier strains showed minimal degradation: *Micrococcus luteus* (12.7%), *Pantoea dispersa* (7.4%), and *Bacillus licheniformis* (3.8%) (Figure 5a). Atrazine degradation patterns revealed different strain leadership, with *P. pergaminensis* again demonstrating superior performance at 62.2% removal (25.70 ± 0.50 ng mL⁻¹ → 9.70 ± 0.10 ng mL⁻¹), followed by *P. polymyxa* (43.1%, 24.10 ± 0.10 ng mL⁻¹ → 13.70 ± 0.20 ng mL⁻¹) and *B. licheniformis* (42.1%, 26.10 ± 0.20 ng mL⁻¹ → 15.10 ± 0.50 ng mL⁻¹). Notably, strains that performed well on PFOS showed markedly reduced atrazine capabilities: *B. cabrialesii* achieved only 11.4% atrazine degradation despite its 40.3% PFOS performance, while *A. hermannii* managed just 6.2% atrazine removal compared to its 32.4% PFOS degradation. Several strains showed negligible atrazine processing: *M. luteus* (6.5%), *P. dispersa* (2.6%), and some strains exhibited concentration increases rather than degradation (Figure 5b). Lignin degradation revealed yet another performance hierarchy, with *P. polymyxa* emerging as the clear leader with 62.7% removal (479.8 ± 23.2 mg L⁻¹ → 178.8 ± 12.1 mg L⁻¹), substantially outperforming its PFOS (18.2%) and atrazine (43.1%) capabilities. *P. pergaminensis* achieved moderate lignin degradation (43.4%, 500.0 ± 14.1 mg L⁻¹ → 283.2 ± 17.4 mg L⁻¹), while *B. pseudomycoides* showed comparable performance (43.0%, 477.8 ± 14.9 mg L⁻¹ → 272.1 ± 15.5 mg L⁻¹). Lower-tier lignin degraders included *B. cabrialesii* (27.9%), *B. licheniformis* (16.7%), *P. fulva* (7.5%), *P. dispersa* (4.6%), and *A. hermannii* (3.4%), with *M. luteus* showing negligible activity (0.6% degradation) (Figure 5c).

**Figure 5.**
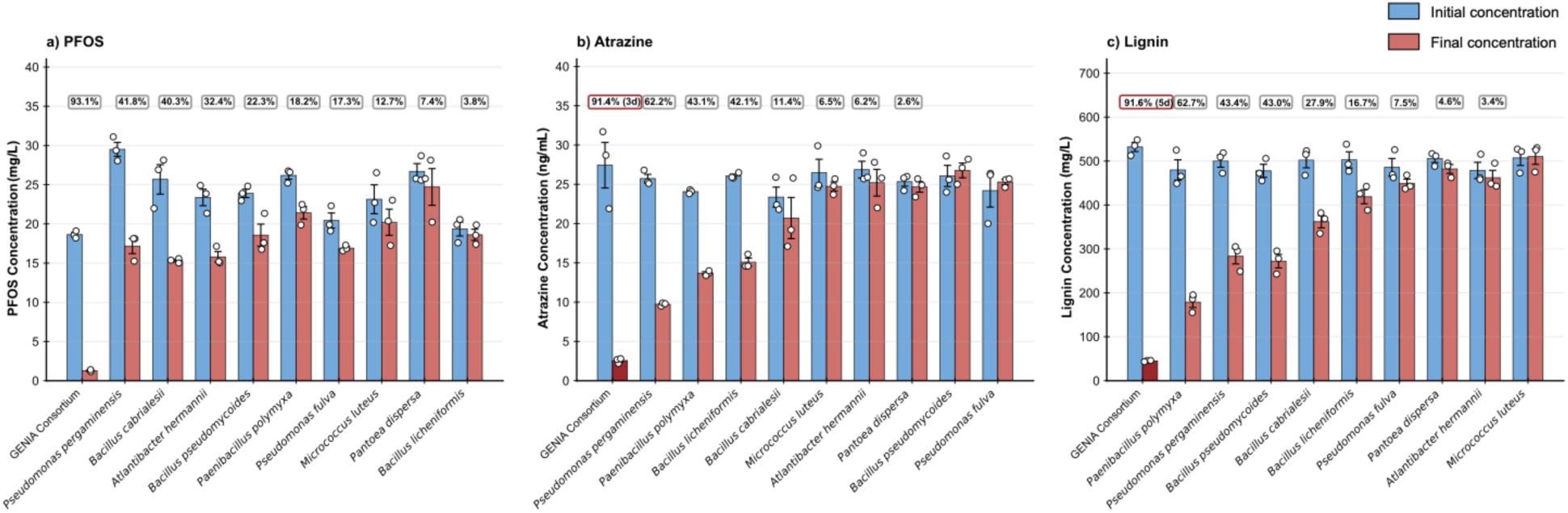
Comparative degradation performance of GENIA consortium versus individual bacterial strains across three pollutant classes. (a) PFOS degradation comparison showing GENIA consortium achieving 93.1% removal efficiency compared to individual strain performance ranging from 3.8% (*Bacillus licheniformis*) to 41.8% (*Pseudomonas pergaminensis*). Blue bars represent initial concentrations, red bars represent final concentrations after 7-day incubation. Individual data points show biological replicates with error bars indicating standard error of the mean. (b) Atrazine degradation performance demonstrating GENIA consortium’s 91.4% removal efficiency achieved in 3 days versus individual strain capabilities ranging from negative values to 62.2% (*P. pergaminensis*) over 7 days. (c) Lignin processing comparison showing GENIA consortium’s 91.6% degradation efficiency in 5 days compared to individual strain performance from 0.6% (*Micrococcus luteus*) to 62.7% (*Paenibacillus polymyxa*) over 7 days. The consortium consistently outperformed the best individual strains across all pollutant classes, demonstrating 2.2-fold improvement in PFOS degradation, 1.5-fold enhancement in atrazine removal, and 1.5-fold increase in lignin processing. All experiments conducted in M9 minimal medium at 27°C with 150 rpm orbital shaking.

### Temporal Lignin Degradation Dynamics

The synthetic microbial consortium demonstrated efficient and comprehensive lignin degradation, with concentrations decreasing from an initial average of 490.7 ± 12.1 mg L⁻¹ to achieve a total removal efficiency of 91.6% by day 5 (Figure 6b). The biodegradation process displayed a distinct temporal progression with sequential concentration reductions: 44.9% elimination within the first day, 82.4% by day 3, 91.6% by day 5, and final concentrations reaching 41.6 ± 1.0 mg L⁻¹, with no substantial variations observed after day 6. This translates to a mean degradation rate of 18.3% daily, demonstrating vigorous metabolic performance against the complex aromatic macromolecule. UV-visible spectral analysis confirmed typical lignin absorption at 287 nm (0.950 ± 0.008 absorbance units) and the generation of hydroxycinnamic acid derivatives at 422 nm through n→π electronic transitions (Fig 6a). Statistical evaluation demonstrated highly significant variations in lignin concentrations throughout the experimental timeframe (one-way ANOVA: p < 0.0001), validating the temporal degradation pattern. The breakdown process exhibited bi-phasic kinetics characterized by rapid initial depolymerization, maintaining consistent enzymatic activity during the entire experimental duration. Simultaneously, hydroxycinnamic acid formation rose from baseline levels (0.019 ± 0.001) to maximum accumulation on day 3 (0.769 ± 0.029), subsequently declining to 0.624 ± 0.019 by day 5, while phenolic intermediate compounds (340 nm) achieved peak concentrations of 0.201 ± 0.008 on day 3 before decreasing to 0.169 ± 0.006 by day 5, substantiating active biodegradation with intermediate metabolite formation and their subsequent biotransformation.

**Figure 6.**
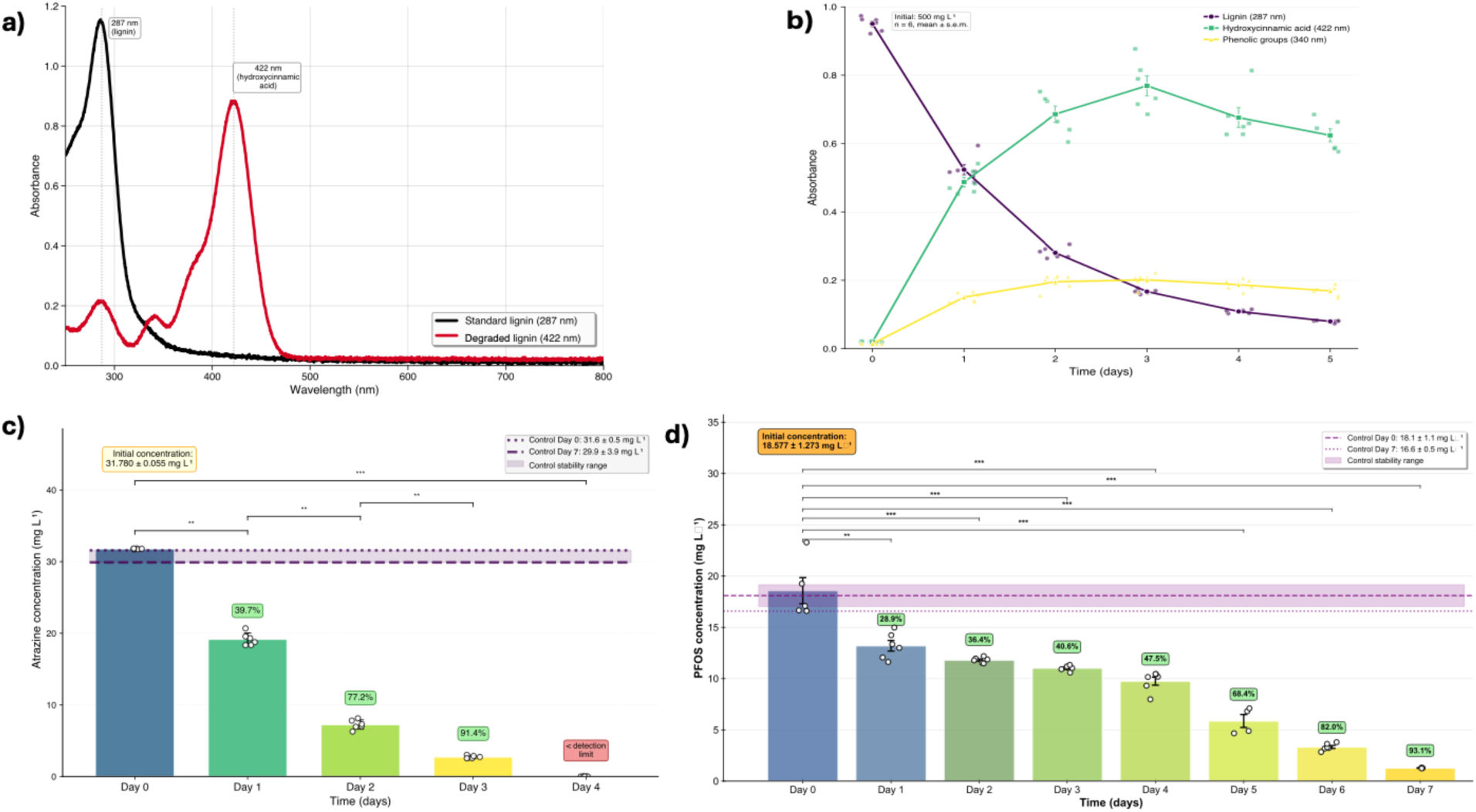
Temporal lignin biodegradation dynamics and intermediate formation by GENIA consortium. (a) UV-Vis spectral analysis showing lignin absorption characteristics at 287 nm (standard lignin, black line) and degradation products at 422 nm (hydroxycinnamic acid derivatives, red line), confirming aromatic compound processing and intermediate formation. (b) Time-course analysis of lignin degradation (287 nm, black line), hydroxycinnamic acid formation (422 nm, green line), and phenolic intermediates (340 nm, yellow line) over 5 days. Initial lignin concentration of 500 mg L⁻¹ declined progressively while intermediate metabolites accumulated to peak concentrations by day 3 before subsequent biotransformation. (c) Atrazine degradation kinetics showing rapid elimination from initial concentration of 31.780 ± 0.055 mg L⁻¹ to below detection limits by day 4, with 91.4% removal achieved by day 3. Control samples (dashed lines) showed no significant degradation, confirming the biologically mediated process. Statistical significance indicated by asterisks (***p < 0.001, **p < 0.01, *p < 0.05). (d) PFOS degradation timeline demonstrating progressive removal from initial concentration of 18.57 ± 2.413 mg L⁻¹ to 93.1% degradation by day 7. Color gradient from blue to yellow indicates temporal progression. Control samples maintained stable concentrations throughout the experimental period. Error bars represent standard error of the mean (n = 3-6 biological replicates per timepoint).

### Temporal Atrazine Degradation Dynamics

The synthetic consortium exhibited rapid and extensive atrazine removal, with concentrations declining from an initial mean of 31.825 ± 0.058 mg L⁻¹ by day 3, representing a total degradation efficiency of 91.4% (Figure 6c). The degradation process followed a clear temporal pattern with progressive concentration reductions: 38.7% removal by day 1 (19.500 ± 0.812 mg L⁻¹), 75.9% by day 2 (7.675 ± 1.204 mg L⁻¹), 91.4% by day 3, and concentrations below detection limits by day 4 (<1 ng mL⁻¹). This corresponds to an average degradation rate of 30.5% per day, indicating robust metabolic activity against the triazine herbicide. Statistical analysis revealed a highly significant difference in atrazine concentrations across the experimental timeline (one-way ANOVA: p < 0.0001), confirming the temporal degradation trend. Linear regression analysis of the concentration decline yielded a strong correlation (R² = 0.969, slope = -9.905 mg L⁻¹ day⁻¹, p = 0.0157), demonstrating consistent first-order degradation kinetics throughout the experimental period. Control treatments maintained stable atrazine concentrations (28.750 ± 4.597 to 31.350 ± 0.661 mg L⁻¹), confirming active biodegradation rather than abiotic removal. Initial concentrations were equivalent between groups (t-test, p = 0.2304), validating the experimental design.

### Temporal PFOS degradation analysis

The synthetic microbial consortium exhibited substantial PFOS degradation capabilities, with concentration declining from an initial mean of 18.58 ± 2.85 mg L⁻¹ to achieve a remarkable removal efficiency of 93.1% by day 7. The biodegradation process demonstrated a progressive temporal pattern with final concentrations reaching 1.28 ± 0.04 mg L⁻¹ by day 7, indicating sustained metabolic activity against this recalcitrant perfluorinated compound. Control samples showed no significant concentration changes (19.61 ± 3.02 mg L⁻¹ at day 0 vs. 16.59 ± 0.95 mg L⁻¹ at day 7; p = 0.173), confirming that observed degradation was biologically mediated rather than due to abiotic processes (Figure 6d). The inverse relationship between PFOS degradation and PFBS accumulation (individual sample correlations ranging from 14.77-17.33 mg L⁻¹ PFOS decrease corresponding to 0.46-1.51 mg L⁻¹ PFBS increase) provides evidence for active defluorination and chain-shortening mechanisms, demonstrating the consortium’s capacity for biotransformation of persistent organofluorine compounds. Concurrent fluoride liberation analysis confirmed active C-F bond cleavage, with fluoride concentrations increasing from undetectable baseline levels following a biphasic kinetic pattern characterized by an initial lag phase through day 1, followed by exponential release beginning on day 2 (2.42 ± 0.32 mg L⁻¹) and reaching final equilibrium of 22.22 ± 0.58 mg L⁻¹ by day 7. The fluoride release exhibited excellent linearity with a rate constant of 4.257 ± 0.333 mg L⁻¹ d⁻¹ (R² = 0.9761, p = 2.16 × 10⁻⁴), providing direct stoichiometric evidence of defluorination activity and confirming that PFOS biotransformation involves systematic fluorine removal rather than simple adsorption or sequestration mechanisms (Supplementary).

## DISCUSSION

As Geoffrey Hinton, the 2024 Nobel Prize in Physics laureate, proclaimed in his acceptance speech: *”This new form of AI excels at modeling human intuition rather than human reasoning, and it will enable us to create highly intelligent and knowledgeable assistants who will increase productivity in almost all industries”* (80). This recognition of artificial intelligence’s transformative potential extends beyond computational applications to fundamental biological systems, where machine learning approaches are revolutionizing our ability to design functional microbial communities for environmental remediation.

### Mechanistic Framework of Synergistic Multi-Pollutant Degradation

Our GENIA-designed consortium operates through a sophisticated metabolic cross-feeding network that fundamentally differs from conventional single-pollutant bioremediation approaches. The >90% PFOS, 91.6% lignin, and 91.4% atrazine removal is attributed to engineered metabolic complementarity where degradation intermediates from one pollutant serve as co-substrates for processing others, creating an integrated biochemical network (81).

The consortium’s lignin degradation pathway exemplifies this synergistic approach. Initial depolymerization by lignin peroxidases (LiP, manganese peroxidases (MnP), and dye-decolorizing peroxidases (DyP) in *Paenibacillus polymyxa* and *Pseudomonas pergaminensis* could yield guaiacyl (G), syringyl (S), and hydroxyphenyl (H) monomers that funnel through vanillate and syringate intermediates (82). These aromatics undergo O-demethylation via LigM and VanAB enzymes to generate protocatechuate and catechol, central intermediates that become available for cross-feeding with other consortium members (83).

For instance, the protocatechuate generated from lignin degradation enters the β-ketoadipate pathway through protocatechuate 3,4-dioxygenase, yielding β-ketoadipate that ultimately produces succinyl-CoA and acetyl-CoA (84). This creates a metabolic bridge where lignin-derived carbon flows could support the energy required for PFAS and atrazine degradation by other consortium members, potentially explaining the enhanced performance compared to strain approaches that achieved only 27.9-67% PFOS degradation (85).

### Quantitative Analysis of Cross-Feeding Dynamics

The enzymatic landscape showed the mechanistic basis for our consortium’s superior performance. Unlike natural microbial communities where metabolite cross-feeding occurs through chance evolutionary optimization (86), our ML-guided design deliberately positions complementary metabolic capabilities to maximize inter-species nutrient exchange. The 22 lignin-degrading enzymes (47.8% of total capacity) positioned across multiple strains ensure continuous production of aromatic intermediates, while 18 PFAS-degrading enzymes (39.1%) and 6 atrazine-degrading enzymes (13.0%) create metabolic sinks that drive the degradation network forward.

Recent studies demonstrate that successful cross-feeding requires specific stoichiometric relationships between metabolite producers and consumers (87). Our GENIA achieves optimal ratios through the hub strain architecture, where *Paenibacillus polymyxa* and *Pseudomonas pergaminensis* function as primary metabolite producers with versatility scores of 3.8 each, supported by specialist consumers that prevent toxic intermediate accumulation. An example of this could be conventional consortia where *Rhodococcus* sp. strain p52 releases catechol during dibenzo-p-dioxin degradation, requiring *Acinetobacter* sp. BD6 as a dedicated detoxification partner (88).

### Multi-Species Optimization Through Metabolic Division of Labor

The consortium’s exceptional performance validates the fundamental principle that engineered multi-species optimization transcends individual strain limitations through strategic metabolic division of labor. This principle is demonstrated by the stark contrast between individual and collective performance: while *Pseudomonas pergaminensis* achieved the highest individual PFOS degradation (41.8%), this represents less than half the consortium’s 93.1% efficiency. Individual strains showed variable and often modest degradation capabilities across different pollutants—*B. cabrialesii* achieved 40.3% PFOS degradation but only 11.4% atrazine removal, while *P. polymyxa* excelled at lignin processing (62.7%) but struggled with PFOS (18.2%)—indicating substrate-specific metabolic constraints that limit single-organism approaches.

The consortium operates through a sophisticated division of labor where each strain assumes leadership for different pollutant classes while contributing complementary functions across the degradation network. *P. pergaminensis* serves as the primary PFAS degrader with its comprehensive haloalkane dehalogenases and cytochrome P450 systems, *P. polymyxa* leads lignin depolymerization through extensive peroxidase networks, and *B. licheniformis* demonstrates specialized atrazine processing capabilities (42.1%). However, the true innovation lies in how strains with moderate individual performance become highly efficient when their specialized functions are coordinated: *B. cabrialesii’s* moderate PFOS degradation (40.3%) combines synergistically with *A. hermannii’s* intermediate processing capabilities (32.4%), while strains like *P. dispersa* provide critical community services through fluoride efflux systems that enable sustained multi-pollutant processing impossible in single-strain systems. This multi-species approach systematically addresses the metabolic bottlenecks that constrain individual organisms: distributed enzymatic capacity prevents pathway saturation, specialized leadership for each pollutant class maximizes degradation efficiency, and cross-feeding networks supply essential cofactors where needed. The quantitative validation of this approach—with 2.2-fold improvement in PFOS degradation, 1.5-fold enhancement in atrazine removal, and 1.5-fold increase in lignin processing compared to best individual performers—demonstrates that coordinated moderate performers collectively achieve superior results than any single high-performing strain, establishing engineered metabolic cooperation as a paradigm-shifting approach where collective metabolic intelligence emerges from rationally orchestrated strain specialization.

### Overcoming the “Forever Chemical” Challenge

The 93.1% PFOS removal achieved by the GENIA-designed consortium represents a breakthrough in PFAS bioremediation, where conventional approaches struggle with recalcitrant C-F bonds that give PFAS their “forever chemical” designation. This exceptional performance could be attributed to distributed defluorination capacity across multiple strains, creating redundant pathways that prevent metabolic bottlenecks (89). The mechanistic pathway may involve initial C-F bond cleavage by haloalkane dehalogenases (DhlA) in *Pseudomonas pergaminensis* and *P. fulva,* generating short-chain perfluorinated intermediates and fluoride ions (90). Direct experimental evidence for this biotransformation pathway was provided by the systematic formation of PFBS (C4) intermediates from PFOS (C8) degradation, with PFBS concentrations increasing from 0.54 mg L⁻¹ on day 6 to 1.40 mg L⁻¹ by day 7, demonstrating active chain-shortening mechanisms that progressively remove fluorinated carbon units. The liberated can be immediately sequestered by CrcB fluoride efflux transporters present in 8/9 consortium strains (88.9% coverage), preventing fluoride toxicity that typically limits PFAS biodegradation, as confirmed by the linear fluoride release kinetics (4.48 mg L⁻¹ final concentration, which means a defluorination percentage of 34.69%). Simultaneously, haloacid dehalogenases (HAD variants) in 4/9 strains process intermediate metabolites, while reduced cofactors (NADPH/NADH) supply the reducing power necessary for continued defluorination via enzymatic electron transfer systems, including NAD(P)H oxidoreductases and cytochrome P450 enzymes.

This distributed approach contrasts with single-strain systems where *Pseudomonas putida* achieved only 19.0% PFOA and 46.9% PFOS transformation in 96 h (85), while *Acidimicrobium* sp. strain A6 demonstrated PFOA degradation with shorter-chain perfluorinated intermediate formation under anaerobic conditions (91). Earlier reports of *Pseudomonas fluorescens* D2 utilizing polyfluorinated H-PFOS as a sulfur source, though unable to transform fully saturated PFOS (92), highlight the metabolic constraints that limit single-organism approaches compared to the designed community’s integrated defluorination network.

### Addressing the Metabolic Bottleneck

The identification of atrazine degradation as the primary metabolic bottleneck (13.0% enzymatic capacity) reflects the complex biochemistry required for triazine ring cleavage. The designed community addresses this through pathway distribution across five strains, employing both hydrolytic and oxidative routes that converge on cyanuric acid as a common intermediate (93). The hydrolytic pathway initiates with atrazine chlorohydrolase (AtzA) in *Pseudomonas pergaminensis* and *Bacillus pseudomycoides*, converting atrazine to hydroxyatrazine through chloride substitution. Sequential dealkylation by AtzB (hydroxyatrazine ethylaminohydrolase) and AtzC (N-isopropylammelide isopropylaminohydrolase) yields cyanuric acid, which undergoes ring cleavage by cyanuric acid hydrolase (AtzE) to produce ammonia and CO₂ (94). Simultaneously, the oxidative pathway employs N-dealkylases and cytochrome P450 enzymes to generate deethylatrazine (DEA) and deisopropylatrazine (DIA) intermediates that ultimately converge on the cyanuric acid node. This pathway redundancy ensures robust atrazine degradation even under varying environmental conditions, explaining the observed 91.4% removal efficiency compared to single-strain approaches that often show variable performance due to nitrogen catabolite repression (95).

The complexity of atrazine metabolism highlighted in our system coincides with a recent work from Zhang et al. (96). This work reported a similar division of labor in a four-member synthetic community achieving 60%−99% degradation efficiency of the endogenous herbicides over 35 days in soil systems, with specialized strains handling different aspects of herbicide metabolism while maintaining functional complementarity.

### Integration with Central Carbon Metabolism

The innovation of the engineered consortium lies in the metabolic integration where degradation products from all three pollutants feed into central carbon metabolism through carefully orchestrated biochemical funneling. Lignin degradation generates protocatechuate and catechol, which undergo ring cleavage to produce β-ketoadipate pathway intermediates including succinyl-CoA, acetyl-CoA, and pyruvate (97). These central metabolites provide the carbon skeletons and energy necessary for biosynthetic processes in low-degrading consortium members. The fluoride generated is not merely detoxified but potentially utilized for specific enzymatic functions in fluoride-tolerant strains. Atrazine degradation yields nitrogen in the form of ammonia, supporting amino acid biosynthesis across the consortium and relieving nitrogen limitation that commonly constrains environmental microbial communities.

### Comparative Performance Analysis

Quantitative comparison with recent literature establishes new benchmarks for multi-pollutant bioremediation. While recent synthetic communities achieved 98.55% imidacloprid degradation in 15 days and 99.33% chlorantraniliprole degradation in 20 days, these studies targeted single pesticides in isolation (96). Comparable studies on lignin degradation show efficiencies reaching 54% after 48 h under optimal conditions (97), while PFAS consortia achieve 56.7% PFOS reduction over 20 days with external co-substrate addition (90). The simultaneous achievement of >90% degradation across all three pollutant classes within 7 days represents a 2-4 fold improvement in efficiency and establishes a complete paradigm shift toward integrated environmental remediation. The temporal kinetics of lignin (bi-phasic, 91.6% by day 5), atrazine (first-order, 91.4% by day 3), and PFOS (93.1% by day 7) demonstrate coordinated metabolic activity that maximizes resource utilization while minimizing inhibitory intermediate accumulation.

### Machine Learning Innovation in Biological System Design

This study represents the first demonstration of machine learning-guided synthetic community assembly for simultaneous degradation of structurally diverse persistent pollutants. The successful application of Graphattention Networks and Node2Vec embeddings (27, 30) to predict optimal microbial community assemblies represents a methodological breakthrough in synthetic biology. Unlike recent applications focused on understanding existing community structure (98) our GENIA framework uniquely targets predictive community design for specific functional outcomes. The 88.9% concordance between GENIA and iNAP 2.0 predictions validates functional relationships that transcend phylogenetic boundaries, representing a significant advance over metabolic modeling approaches requiring extensive manual curation (99, 100). Our framework automates metabolic complementarity identification and predicts community stability under perturbation, enabling rational biological system design with unprecedented precision.

### Environmental and Economic Implications

The demonstration of simultaneous multi-pollutant degradation addresses critical gaps in environmental biotechnology where contaminated sites contain complex pollutant mixtures requiring costly sequential treatments (101). As regulatory pressure increases for comprehensive environmental remediation, particularly for persistent pollutants like PFAS (8), engineered communities with defined functional capabilities become increasingly valuable for both remediation and prevention strategies. The framework’s modularity presented here enables expansion to additional pollutant classes, providing a scalable platform for addressing emerging contaminant challenges (93).

The GENIA framework establishes a proof-of-concept for machine learning-guided design of synthetic microbial communities, demonstrating for the first time that computational approaches can successfully predict and engineer biological systems for complex environmental applications. As machine learning continues to revolutionize biological sciences, our approach exemplifies how artificial intelligence can capture biological intuition to design functional microbial consortia. These consortia possess emergent properties exceeding those of individual strains or natural communities, opening new avenues for addressing global environmental challenges through engineered biological solutions.

## ACKNOWLEDGEMENTS

This research was supported by the United States Department of Commerce National Institute of Standards and Technology Award (60NANB24D118). Authors declare no conflict of interest.

## AUTHOR CONTRIBUTIONS

Esaú De la Vega-Camarillo, Jorge Arreola-Vargas, Sanjay Antony-Babu and Won Bo Shim conceived and designed the experiments. Esaú De la Vega-Camarillo, Saurav Kumar Mathur, and Joshua Andrew Santos performed experiments and analyses. Esaú De la Vega-Camarillo, Jorge Arreola-Vargas, Sanjay Antony-Babu and Won Bo Shim wrote the manuscript.

## REFERENCES

1. Margulis, L. Symbiosis in Cell Evolution: Life and Its Environment on the Early Earth; W. H. Freeman: San Francisco, 1981.

2. Lindemann, S. R.; Bernstein, H. C.; Song, H.-S.; Fredrickson, J. K.; Fields, M. W.; Shou, W.; Johnson, D. R.; Beliaev, A. S. Engineering Microbial Consortia for Controllable Outputs. The ISME Journal 2016, 10 (9), 2077–2084. 10.1038/ismej.2016.26.

3. McCarty, N. S.; Ledesma-Amaro, R. Synthetic Biology Tools to Engineer Microbial Communities for Biotechnology. Trends in Biotechnology 2019, 37 (2), 181–197. 10.1016/j.tibtech.2018.11.002.

4. Shetty, S. A.; Hugenholtz, F.; Lahti, L.; Smidt, H.; De Vos, W. M. Intestinal Microbiome Landscaping: Insight in Community Assemblage and Implications for Microbial Modulation Strategies. FEMS Microbiology Reviews 2017, 41 (2), 182–199. 10.1093/femsre/fuw045.

5. Ghattas, A.-K.; Fischer, F.; Wick, A.; Ternes, T. A. Anaerobic Biodegradation of (Emerging) Organic Contaminants in the Aquatic Environment. Water Research 2017, 116, 268–295. 10.1016/j.watres.2017.02.001.

6. Khatoon, H.; Rai, J. P. N. Optimization Studies on Biodegradation of Atrazine by Bacillus Badius ABP6 Strain Using Response Surface Methodology. Biotechnology Reports 2020, 26, e00459. 10.1016/j.btre.2020.e00459.

7. Ahmad, M.; Riaz, U.; Iqbal, S.; Ahmad, J.; Rasheed, H.; Al-Farraj, A. S. F.; Al-Wabel, M. I. Adsorptive Removal of Atrazine From Contaminated Water Using Low-Cost Carbonaceous Materials: A Review. Front. Mater. 2022, 9. 10.3389/fmats.2022.909534.

8. Cousins, I. T.; DeWitt, J. C.; Glüge, J.; Goldenman, G.; Herzke, D.; Lohmann, R.; Ng, C. A.; Scheringer, M.; Wang, Z. The High Persistence of PFAS Is Sufficient for Their Management as a Chemical Class. Environ. Sci.: Processes Impacts 2020, 22 (12), 2307–2312. 10.1039/d0em00355g.

9. Hayes, T. B.; Collins, A.; Lee, M.; Mendoza, M.; Noriega, N.; Stuart, A. A.; Vonk, A. Hermaphroditic, Demasculinized Frogs after Exposure to the Herbicide Atrazine at Low Ecologically Relevant Doses. Proc. Natl. Acad. Sci. U.S.A. 2002, 99 (8), 5476–5480. 10.1073/pnas.082121499.

10. Qu, M.; Liu, G.; Zhao, J.; Li, H.; Liu, W.; Yan, Y.; Feng, X.; Zhu, D. Fate of Atrazine and Its Relationship with Environmental Factors in Distinctly Different Lake Sediments Associated with Hydrophytes. Environmental Pollution 2020, 256, 113371. 10.1016/j.envpol.2019.113371.

11. Wang, Y.; Lin, T.; Chen, H. Degradation of Atrazine by a UV-Activated Organic Chloramines Process: Kinetics, Degradation Pathways, Disinfection by-Product Formation, and Toxicity Changes. Chemical Engineering Journal 2023, 468, 143788. 10.1016/j.cej.2023.143788.

12. Zakzeski, J.; Bruijnincx, P. C. A.; Jongerius, A. L.; Weckhuysen, B. M. The Catalytic Valorization of Lignin for the Production of Renewable Chemicals. Chem. Rev. 2010, 110 (6), 3552–3599. 10.1021/cr900354u.

13. Bugg, T. D.; Ahmad, M.; Hardiman, E. M.; Singh, R. The Emerging Role for Bacteria in Lignin Degradation and Bio-Product Formation. Current Opinion in Biotechnology 2011, 22 (3), 394–400. 10.1016/j.copbio.2010.10.009.

14. Renner, R. Growing Concern Over Perfluorinated Chemicals. Environ. Sci. Technol. 2001, 35 (7), 154A–160A. 10.1021/es012317k.

15. D’Agostino, L. A.; Mabury, S. A. Identification of Novel Fluorinated Surfactants in Aqueous Film Forming Foams and Commercial Surfactant Concentrates. Environ. Sci. Technol. 2014, 48 (1), 121–129. 10.1021/es403729e.

16. Dasu, K.; Liu, J.; Lee, L. S. Aerobic Soil Biodegradation of 8:2 Fluorotelomer Stearate Monoester. Environ. Sci. Technol. 2012, 46 (7), 3831–3836. 10.1021/es203978g.

17. Harris, B. A.; Zhou, J.; Clarke, B. O.; Leung, I. K. H. Enzymatic Degradation of PFAS: Current Status and Ongoing Challenges. ChemSusChem 2025, 18 (2). 10.1002/cssc.202401122.

18. Henn, C.; Monteiro, D. A.; Boscolo, M.; Da Silva, R.; Gomes, E. Biodegradation of Atrazine and Ligninolytic Enzyme Production by Basidiomycete Strains. BMC Microbiol 2020, 20 (1). 10.1186/s12866-020-01950-0.

19. Huang, S.; Jaffé, P. R. Defluorination of Perfluorooctanoic Acid (PFOA) and Perfluorooctane Sulfonate (PFOS) by *Acidimicrobium* Sp. Strain A6. Environ. Sci. Technol. 2019, 53 (19), 11410–11419. 10.1021/acs.est.9b04047.

20. Lozupone, C. A.; Knight, R. Global Patterns in Bacterial Diversity. Proc. Natl. Acad. Sci. U.S.A. 2007, 104 (27), 11436–11440. 10.1073/pnas.0611525104.

21. Díaz-Rodríguez, A. M.; Salcedo Gastelum, L. A.; Félix Pablos, C. M.; Parra-Cota, F. I.; Santoyo, G.; Puente, M. L.; Bhattacharya, D.; Mukherjee, J.; De Los Santos-Villalobos, S. The Current and Future Role of Microbial Culture Collections in Food Security Worldwide. Front. Sustain. Food Syst. 2021, 4. 10.3389/fsufs.2020.614739.

22. Liang, J.; Arellano, A.; Sajjadi, S.; Jewell, T. Cultivation of Diverse and Rare Bacteria from the Human Gut Microbiome: Rapid Isolation of Entire Populations of Microbes Using the GALT Prospector, an Automated Array-Based Platform. Genetic Engineering & Biotechnology News 2020, 40 (7), 50–52. 10.1089/gen.40.07.14.

23. Nichols, D.; Cahoon, N.; Trakhtenberg, E. M.; Pham, L.; Mehta, A.; Belanger, A.; Kanigan, T.; Lewis, K.; Epstein, S. S. Use of Ichip for High-Throughput *In Situ* Cultivation of “Uncultivable” Microbial Species. Appl Environ Microbiol 2010, 76 (8), 2445–2450. 10.1128/aem.01754-09.

24. Simpson, J. T.; Pop, M. The Theory and Practice of Genome Sequence Assembly. Annu. Rev. Genom. Hum. Genet. 2015, 16 (1), 153–172. 10.1146/annurev-genom-090314-050032.

25. Wick, R. R.; Judd, L. M.; Gorrie, C. L.; Holt, K. E. Completing Bacterial Genome Assemblies with Multiplex MinION Sequencing. Microbial Genomics 2017, 3 (10). 10.1099/mgen.0.000132.

26. McGann, P.; Bunin, J. L.; Snesrud, E.; Singh, S.; Maybank, R.; Ong, A. C.; Kwak, Y. I.; Seronello, S.; Clifford, R. J.; Hinkle, M.; Yamada, S.; Barnhill, J.; Lesho, E. Real Time Application of Whole Genome Sequencing for Outbreak Investigation – What Is an Achievable Turnaround Time? Diagnostic Microbiology and Infectious Disease 2016, 85 (3), 277–282. 10.1016/j.diagmicrobio.2016.04.020.

27. Valous, N. A.; Popp, F.; Zörnig, I.; Jäger, D.; Charoentong, P. Graph Machine Learning for Integrated Multi-Omics Analysis. Br J Cancer 2024, 131 (2), 205–211. 10.1038/s41416-024-02706-7.

28. Zitnik, M.; Nguyen, F.; Wang, B.; Leskovec, J.; Goldenberg, A.; Hoffman, M. M. Machine Learning for Integrating Data in Biology and Medicine: Principles, Practice, and Opportunities. Information Fusion 2019, 50, 71–91. 10.1016/j.inffus.2018.09.012.

29. Dornaika, F.; Bi, J.; Charafeddine, J. Leveraging Graph Convolutional Networks for Semi-Supervised Learning in Multi-View Non-Graph Data. Cogn Comput 2025, 17 (2). 10.1007/s12559-025-10428-y.

30. Grover, A.; Leskovec, J. Node2vec: Scalable Feature Learning for Networks. In Proceedings of the 22nd ACM SIGKDD International Conference on Knowledge Discovery and Data Mining; ACM: San Francisco California USA, 2016; pp 855–864. 10.1145/2939672.2939754.

31. Hamilton, W. L.; Ying, R.; Leskovec, J. Inductive Representation Learning on Large Graphs. arXiv September 10, 2018. 10.48550/arXiv.1706.02216.

32. Alluhaybi, A. A.; Alharbi, A.; Alshammari, K. F.; El-Desouky, M. G. Efficient Adsorption and Removal of the Herbicide 2,4-Dichlorophenylacetic Acid from Aqueous Solutions Using MIL-88(Fe)-NH_2_. ACS Omega 2023, 8 (43), 40775–40784. 10.1021/acsomega.3c05818.

33. Sagarkar, S.; Mukherjee, S.; Nousiainen, A.; Björklöf, K.; Purohit, H. J.; Jørgensen, K. S.; Kapley, A. Monitoring Bioremediation of Atrazine in Soil Microcosms Using Molecular Tools. Environmental Pollution 2013, 172, 108–115. 10.1016/j.envpol.2012.07.048.

34. Isolation Bio. Prospector® Microbial Isolation and Cultivation System: User Manual, Version 1.2; Isolation Bio, Inc.: San Carlos, CA, 2023.

35. Persyn, A.; Mueller, A.; Goormachtig, S. Drops Join to Make a Stream: High-throughput Nanoscale Cultivation to Grasp the Lettuce Root Microbiome. Environ Microbiol Rep 2022, 14 (1), 60–69. 10.1111/1758-2229.13014.

36. Opentrons OT-2 Liquid Handling Platform. Opentrons Labworks Inc., Brooklyn, NY. https://opentrons.com/products/ot-2-robot (accessed July 12, 2025).

37. Boundy-Mills, K. L.; de Jonge, R.; Flaherty, J. E.; Giunta, A.; Heckman, D. Mechanisms of Atrazine Biodegradation in Agricultural Soils. Appl. Environ. Microbiol. 2017, 83 (18), e00816–17. DOI: 10.1128/AEM.00816-17.

38. Liu, J.; Mejia Avendaño, S. Microbial Degradation of Polyfluoroalkyl Chemicals in the Environment: A Review. Environ. Int. 2013, 61, 98–114. DOI: 10.1016/j.envint.2013.08.022.

39. Kolmogorov, M.; Yuan, J.; Lin, Y.; Pevzner, P. Assembly of Long Error-Prone Reads Using Repeat Graphs. Nat. Biotechnol. 2019, 37 (5), 540–546. DOI: 10.1038/s41587-019-0072-8.

40. Vaser, R.; Sović, I.; Nagarajan, N.; Šikić, M. Fast and Accurate De Novo Genome Assembly from Long Uncorrected Reads. Genome Res. 2017, 27 (5), 737–746. DOI: 10.1101/gr.214270.116.

41. Oxford Nanopore Technologies. Medaka: Sequence Correction Provided by ONT Research. Oxford, UK. https://github.com/nanoporetech/medaka (accessed July 12, 2025).

42. Gurevich, A.; Saveliev, V.; Vyahhi, N.; Tesler, G. QUAST: Quality Assessment Tool for Genome Assemblies. Bioinformatics 2013, 29 (8), 1072–1075. DOI: 10.1093/bioinformatics/btt086.

43. Parks, D. H.; Imelfort, M.; Skennerton, C. T.; Hugenholtz, P.; Tyson, G. W. CheckM: Assessing the Quality of Microbial Genomes Recovered from Isolates, Single Cells, and Metagenomes. Genome Res. 2015, 25 (7), 1043–1055. DOI: 10.1101/gr.186072.114.

44. Tatusova, T.; DiCuccio, M.; Badretdin, A.; Chetvernin, V.; Nawrocki, E. P.; Zaslavsky, L.; Lomsadze, A.; Pruitt, K. D.; Borodovsky, M.; Ostell, J. NCBI Prokaryotic Genome Annotation Pipeline. Nucleic Acids Res. 2016, 44 (14), 6614–6624. DOI: 10.1093/nar/gkw569.

45. Seemann, T. Prokka: Rapid Prokaryotic Genome Annotation. Bioinformatics 2014, 30 (14), 2068–2069. DOI: 10.1093/bioinformatics/btu153.

46. Aziz, R. K.; Bartels, D.; Best, A. A.; DeJongh, M.; Disz, T.; Edwards, R. A.; Formsma, K.; Gerdes, S.; Glass, E. M.; Kubal, M.; Meyer, F.; Olsen, G. J.; Olson, R.; Osterman, A. L.; Overbeek, R. A.; McNeil, L. K.; Paarmann, D.; Paczian, T.; Parrello, B.; Pusch, G. D.; Reich, C.; Stevens, R.; Vassieva, O.; Vonstein, V.; Wilke, A.; Zagnitko, O. The RAST Server: Rapid Annotations Using Subsystems Technology. BMC Genomics 2008, 9, 75. DOI: 10.1186/1471-2164-9-75.

47. Hyatt, D.; Chen, G.-L.; LoCascio, P. F.; Land, M. L.; Larimer, F. W.; Hauser, L. J. Prodigal: Prokaryotic Gene Recognition and Translation Initiation Site Identification. BMC Bioinformatics 2010, 11, 119. DOI: 10.1186/1471-2105-11-119.

48. Kanehisa, M.; Goto, S. KEGG: Kyoto Encyclopedia of Genes and Genomes. Nucleic Acids Res. 2000, 28 (1), 27–30. DOI: 10.1093/nar/28.1.27.

49. Galperin, M. Y.; Makarova, K. S.; Wolf, Y. I.; Koonin, E. V. Expanded Microbial Genome Coverage and Improved Protein Family Annotation in the COG Database. Nucleic Acids Res. 2015, 43 (D1), D261–D269. DOI: 10.1093/nar/gku1223.

50. King, Z. A.; Lu, J.; Dräger, A.; Miller, P.; Federowicz, S.; Lerman, J. A.; Ebrahim, A.; Palsson, B. O.; Lewis, N. E. BiGG Models: A Platform for Integrating, Standardizing and Sharing Genome-Scale Models. Nucleic Acids Res. 2016, 44 (D1), D515–D522. DOI: 10.1093/nar/gkv1049.

51. Altschul, S. F.; Gish, W.; Miller, W.; Myers, E. W.; Lipman, D. J. Basic Local Alignment Search Tool. J. Mol. Biol. 1990, 215 (3), 403–410. DOI: 10.1016/S0022-2836(05)80360-2.

52. Avsec, Ž.; Weilert, M.; Shrikumar, A.; Krueger, S.; Alexandari, A.; Dalal, K.; Fropf, R.; McAnany, C.; Gagneur, J.; Kundaje, A.; Zeitlinger, J. Modeling Positional Effects of Regulatory Sequences with Spline Transformations Increases Prediction Accuracy of Deep Neural Networks. Bioinformatics 2018, 34 (8), 1261–1268. DOI: 10.1093/bioinformatics/btw727.

53. Karp, P. D.; Weaver, D.; Latendresse, M. How Accurate Is Automated Gap Filling of Metabolic Models? BMC Syst. Biol. 2018, 12 (1), 73. DOI: 10.1186/s12918-018-0593-7.

54. Reed, J. L.; Vo, T. D.; Schilling, C. H.; Palsson, B. O. An Expanded Genome-Scale Model of Escherichia coli K-12 (iJR904 GSM/GPR). Genome Biol. 2003, 4 (9), R54. DOI: 10.1186/gb-2003-4-9-r54.

55. Burgard, A. P.; Nikolaev, E. V.; Schilling, C. H.; Maranas, C. D. Flux Coupling Analysis of Genome-Scale Metabolic Network Reconstructions. Genome Res. 2004, 14 (2), 301–312. DOI: 10.1101/gr.1926504.

56. Zenil, H.; Kiani, N. A.; Tegnér, J. Methods of Information Theory and Algorithmic Complexity for Network Biology. Semin. Cell Dev. Biol. 2016, 51, 32–43. DOI: 10.1016/j.semcdb.2016.01.011.

57. Li, M.; Vitányi, P. An Introduction to Kolmogorov Complexity and Its Applications, 3rd ed.; Springer: New York, 2008.

58. Mahadevan, R.; Edwards, J. S.; Doyle, F. J. Dynamic Flux Balance Analysis of Diauxic Growth in Escherichia coli. Biophys. J. 2002, 83 (3), 1331–1340. DOI: 10.1016/S0006-3495(02)73903-9.

59. Zhuang, K.; Izallalen, M.; Mouser, P.; Richter, H.; Risso, C.; Mahadevan, R.; Lovley, D. R. Genome-Scale Dynamic Modeling of the Competition between Rhodoferax and Geobacter in Anoxic Subsurface Environments. ISME J. 2011, 5 (2), 305–316. DOI: 10.1038/ismej.2010.117.

60. Veličković, P.; Cucurull, G.; Casanova, A.; Romero, A.; Liò, P.; Bengio, Y. Graph Attention Networks. arXiv preprint 2017, arXiv:1710.10903. DOI: 10.48550/arXiv.1710.10903.

61. Grover, A.; Leskovec, J. node2vec: Scalable Feature Learning for Networks. In Proceedings of the 22nd ACM SIGKDD International Conference on Knowledge Discovery and Data Mining; ACM: New York, 2016; pp 855–864. DOI: 10.1145/2939672.2939754.

62. Peng, X.; Feng, K.; Yang, X.; He, Q.; Zhao, B.; Li, T.; Wang, S.; Deng, Y. iNAP 2.0: Harnessing Metabolic Complementarity in Microbial Network Analysis. iMeta 2024, 3 (4), e235. DOI: 10.1002/imt2.235.

63. Mirzaei, R.; Akbari, A.; Shayestehpour, M.; Jafari, M.; Shahsavari, E. Multi-Residue Analysis of Herbicides in Environmental Waters by Liquid Chromatography-Tandem Mass Spectrometry. J. Chromatogr. A 2019, 1608, 460415. DOI: 10.1016/j.chroma.2019.460415.

64. Fujii, S.; Polprasert, C.; Tanaka, S.; Lien, N. P. H.; Qiu, Y. New POPs in the Water Environment: Distribution, Bioaccumulation and Treatment of Perfluorinated Compounds - A Review Paper. J. Water Supply Res. Technol. 2007, 56 (5), 313–326. DOI: 10.2166/aqua.2007.005.

65. Hach Company. SPADNS 2 Method 10225: Fluoride Determination Using Accuvac® Ampules. EPA Method SM 4500-F D Equivalent; Hach Company: Loveland, CO, 2021.

66. Lee, R. A.; Bédard, C.; Berberi, V.; Beauchet, R.; Lavoie, J.-M. UV–Vis as Quantification Tool for Solubilized Lignin Following a Single-Shot Steam Process. Bioresource Technology 2013, 144, 658–663. 10.1016/j.biortech.2013.06.045.

67. Abadi, M.; Agarwal, A.; Barham, P.; Brevdo, E.; Chen, Z.; Citro, C.; Corrado, G. S.; Davis, A.; Dean, J.; Devin, M.; et al. TensorFlow: Large-Scale Machine Learning on Heterogeneous Distributed Systems. arXiv preprint 2016, arXiv:1603.04467.

68. Paszke, A.; Gross, S.; Massa, F.; Lerer, A.; Bradbury, J.; Chanan, G.; Killeen, T.; Lin, Z.; Gimelshein, N.; Antiga, L.; et al. PyTorch: An Imperative Style, High-Performance Deep Learning Library. In Advances in Neural Information Processing Systems 32; Curran Associates, Inc.: 2019; pp 8024–8035.

69. Pedregosa, F.; Varoquaux, G.; Gramfort, A.; Michel, V.; Thirion, B.; Grisel, O.; Blondel, M.; Prettenhofer, P.; Weiss, R.; Dubourg, V.;, et al. Scikit-learn: Machine Learning in Python. J. Mach. Learn. Res. 2011, 12, 2825–2830.

70. Harris, C. R.; Millman, K. J.; van der Walt, S. J.; Gommers, R.; Virtanen, P.; Cournapeau, D.; Wieser, E.; Taylor, J.; Berg, S.; Smith, N. J.;, et al. Array Programming with NumPy. Nature 2020, 585 (7825), 357–362. DOI: 10.1038/s41586-020-2649-2.

71. Chen, J.; Bittinger, K.; Charlson, E. S.; Hoffmann, C.; Lewis, J.; Wu, G. D.; Collman, R. G.; Bushman, F. D.; Li, H. Associating Microbiome Composition with Environmental Covariates Using Generalized UniFrac Distances. Bioinformatics 2012, 28 (16), 2106–2113. DOI: 10.1093/bioinformatics/bts342.

72. Zhao, N.; Chen, J.; Carroll, I. M.; Ringel-Kulka, T.; Epstein, M. P.; Zhou, H.; Zhou, J. J.; Ringel, Y.; Li, H.; Wu, M. C. Testing in Microbiome-Profiling Studies with MiRKAT, the Microbiome Regression-Based Kernel Association Test. Am. J. Hum. Genet. 2015, 96 (5), 797–807. DOI: 10.1016/j.ajhg.2015.04.003.

73. Koh, H.; Livanos, A. E.; Blaser, M. J.; Li, H. A Powerful Microbial Group Association Test Based on the Higher Criticism Analysis for Sparse Microbial Association Signals. Microbiome 2020, 8 (1), 63. DOI: 10.1186/s40168-020-00834-9.

74. Anderson, M. J. Permutational Multivariate Analysis of Variance (PERMANOVA). In Wiley StatsRef: Statistics Reference Online; 2017; pp 1–15. DOI: 10.1002/9781118445112.stat07841.

75. Akiba, T.; Sano, S.; Yanase, T.; Ohta, T.; Koyama, M. Optuna: A Next-Generation Hyperparameter Optimization Framework. In Proceedings of the 25th ACM SIGKDD International Conference on Knowledge Discovery & Data Mining; ACM: 2019; pp 2623–2631. DOI: 10.1145/3292500.3330701.

76. Loshchilov, I.; Hutter, F. SGDR: Stochastic Gradient Descent with Warm Restarts. arXiv May 3, 2017. 10.48550/arXiv.1608.03983.

77. Pascanu, R.; Mikolov, T.; Bengio, Y. On the Difficulty of Training Recurrent Neural Networks. arXiv February 16, 2013. 10.48550/arXiv.1211.5063.

78. Lundberg, S.; Lee, S.-I. A Unified Approach to Interpreting Model Predictions. arXiv November 25, 2017. 10.48550/arXiv.1705.07874.

79. Sundararajan, M.; Taly, A.; Yan, Q. Axiomatic Attribution for Deep Networks. arXiv June 13, 2017. 10.48550/arXiv.1703.01365.

80. Hinton, G. E. Nobel Lecture: Physics 2024. NobelPrize.org. https://www.nobelprize.org/prizes/physics/2024/hinton/speech/ (accessed July 28, 2025).

81. Fritts, R. K.; McCully, A. L.; McKinlay, J. B. Extracellular Metabolism Sets the Table for Microbial Cross-Feeding. Microbiol Mol Biol Rev 2021, 85 (1). 10.1128/mmbr.00135-20.

82. Xu, Z.; Lei, P.; Zhai, R.; Wen, Z.; Jin, M. Recent Advances in Lignin Valorization with Bacterial Cultures: Microorganisms, Metabolic Pathways, and Bio-Products. Biotechnol Biofuels 2019, 12 (1). 10.1186/s13068-019-1376-0.

83. Gu, J.; Qiu, Q.; Yu, Y.; Sun, X.; Tian, K.; Chang, M.; Wang, Y.; Zhang, F.; Huo, H. Bacterial Transformation of Lignin: Key Enzymes and High-Value Products. Biotechnol Biofuels 2024, 17 (1). 10.1186/s13068-023-02447-4.

84. Dos Santos Melo-Nascimento, A. O.; Mota Moitinho Sant’Anna, B.; Gonçalves, C. C.; Santos, G.; Noronha, E.; Parachin, N.; De Abreu Roque, M. R.; Bruce, T. Complete Genome Reveals Genetic Repertoire and Potential Metabolic Strategies Involved in Lignin Degradation by Environmental Ligninolytic Klebsiella Variicola P1CD1. PLoS ONE 2020, 15 (12), e0243739. 10.1371/journal.pone.0243739.

85. Chiriac, F. L.; Stoica, C.; Iftode, C.; Pirvu, F.; Petre, V. A.; Paun, I.; Pascu, L. F.; Vasile, G. G.; Nita-Lazar, M. Bacterial Biodegradation of Perfluorooctanoic Acid (PFOA) and Perfluorosulfonic Acid (PFOS) Using Pure Pseudomonas Strains. Sustainability 2023, 15 (18), 14000. 10.3390/su151814000.

86. Ge, Z.-B.; Zhai, Z.-Q.; Xie, W.-Y.; Dai, J.; Huang, K.; Johnson, D. R.; Zhao, F.-J.; Wang, P. Two-Tiered Mutualism Improves Survival and Competitiveness of Cross-Feeding Soil Bacteria. The ISME Journal 2023, 17 (11), 2090–2102. 10.1038/s41396-023-01519-5.

87. Liao, C.; Wang, T.; Maslov, S.; Xavier, J. B. Modeling Microbial Cross-Feeding at Intermediate Scale Portrays Community Dynamics and Species Coexistence. PLoS Comput Biol 2020, 16 (8), e1008135. 10.1371/journal.pcbi.1008135.

88. Zhao, W.; Wu, Y.; Fu, C.; Li, L. Mutualistic Cross Feeding Mediated by Metabolic Intermediates and Siderophores Enhances Dibenzo-*p*-Dioxin Removal. Environ. Sci. Technol. 2025, 59 (28), 14508–14517. 10.1021/acs.est.4c11857.

89. Harris, B. A.; Zhou, J.; Clarke, B. O.; Leung, I. K. H. Enzymatic Degradation of PFAS: Current Status and Ongoing Challenges. ChemSusChem 2025, 18 (2). 10.1002/cssc.202401122.

90. Liang, Y.; Ma, A. Investigating the Degradation Potential of Microbial Consortia for Perfluorooctane Sulfonate through a Functional “Top-down” Screening Approach. PLoS ONE 2024, 19 (5), e0303904. 10.1371/journal.pone.0303904.

91. Skinner, J. P.; Raderstorf, A.; Rittmann, B. E.; Delgado, A. G. Biotransforming the “Forever Chemicals”: Trends and Insights from Microbiological Studies on PFAS. Environ. Sci. Technol. 2025, 59 (11), 5417–5430. 10.1021/acs.est.4c04557.

92. LaFond, J. A.; Hatzinger, P. B.; Guelfo, J. L.; Millerick, K.; Jackson, W. A. Bacterial Transformation of Per- and Poly-Fluoroalkyl Substances: A Review for the Field of Bioremediation. Environ. Sci.: Adv. 2023, 2 (8), 1019–1041. 10.1039/d3va00031a.

93. Shah, B. A.; Malhotra, H.; Papade, S. E.; Dhamale, T.; Ingale, O. P.; Kasarlawar, S. T.; Phale, P. S. Microbial Degradation of Contaminants of Emerging Concern: Metabolic, Genetic and Omics Insights for Enhanced Bioremediation. Front. Bioeng. Biotechnol. 2024, 12. 10.3389/fbioe.2024.1470522.

94. Govantes, F.; Porrúa, O.; García-González, V.; Santero, E. Atrazine Biodegradation in the Lab and in the Field: Enzymatic Activities and Gene Regulation. Microbial Biotechnology 2009, 2 (2), 178–185. 10.1111/j.1751-7915.2008.00073.x.

95. Zhao, X.; Wang, L.; Ma, F.; Yang, J. Characterisation of an Efficient Atrazine-Degrading Bacterium, Arthrobacter Sp. ZXY-2: An Attempt to Lay the Foundation for Potential Bioaugmentation Applications. Biotechnol Biofuels 2018, 11 (1). 10.1186/s13068-018-1113-0.

96. Zhang, Y.; Gilbert, J. A.; Liu, X.; Nie, L.; Xu, X.; Gao, G.; Lyu, L.; Ma, Y.; Fan, K.; Yang, T.; Zhang, Y.; Zhang, J.; Chu, H. SynCom-mediated Herbicide Degradation Activates Microbial Carbon Metabolism in Soils. iMeta 2025. 10.1002/imt2.70058.

97. Zhang, R.; Wang, J.; Milligan, S.; Yan, Y. Microbial Utilization of Lignin-Derived Aromatics *via* a Synthetic Catechol *Meta*-Cleavage Pathway. Green Chem. 2021, 23 (20), 8238–8250. 10.1039/d1gc02347k.

98. Przymus, P.; Rykaczewski, K.; Martín-Segura, A.; Truu, J.; Carrillo De Santa Pau, E.; Kolev, M.; Naskinova, I.; Gruca, A.; Sampri, A.; Frohme, M.; Nechyporenko, A. Deep Learning in Microbiome Analysis: A Comprehensive Review of Neural Network Models. Front. Microbiol. 2025, 15. 10.3389/fmicb.2024.1516667.

99. Karp, P. D.; Weaver, D.; Latendresse, M. How Accurate Is Automated Gap Filling of Metabolic Models? BMC Syst Biol 2018, 12 (1). 10.1186/s12918-018-0593-7.

100. Silva-Andrade, C.; Rodriguez-Fernández, M.; Garrido, D.; Martin, A. J. M. Using Metabolic Networks to Predict Cross-Feeding and Competition Interactions between Microorganisms. Microbiol Spectr 2024, 12 (5). 10.1128/spectrum.02287-23.

101. Kim, S.-B.; Lee, S.-Y.; Park, S.-J. TiO2/Multi-Walled Carbon Nanotube Electrospun Nanofibers Mats for Enhanced Cr(VI) Photoreduction. Journal of Cleaner Production 2024, 448, 141611. 10.1016/j.jclepro.2024.141611.

