## Supplementary figures for "Machine Learning-Guided Synthetic Microbial Communities Enable Functional and Sustainable Degradation of Persistent Environmental Pollutants"

**SUPPLEMENTARY INFORMATION**

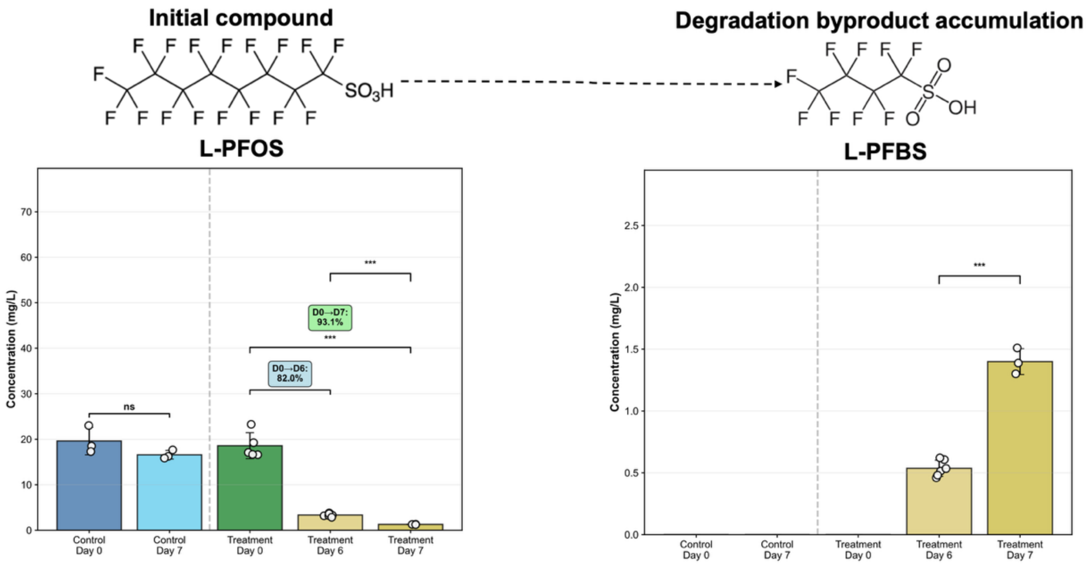


**Figure S1. PFOS biodegradation and chain-shortening mechanism demonstrated by the GENIA consortium.** The figure shows the biotransformation pathway from PFOS (perfluorooctane sulfonate, C8) to PFBS (perfluorobutane sulfonate, C4) through systematic chain-shortening. The left panel displays PFOS concentration decline over 7 days, with significant degradation occurring from day 6 (82.0% removal) to day 7 (93.1% removal) compared to stable control concentrations. The right panel shows corresponding PFBS accumulation as the primary degradation byproduct, with concentrations increasing from undetectable levels at day 0 to $0.54 \pm 0.05$ mg $L^{-1}$ at day 6 and $1.40 \pm 0.04$ mg $L^{-1}$ at day 7. Statistical significance indicated by asterisks (***$p < 0.001$, ns = not significant). Error bars represent the standard error of the mean (n = 3-6 biological replicates).

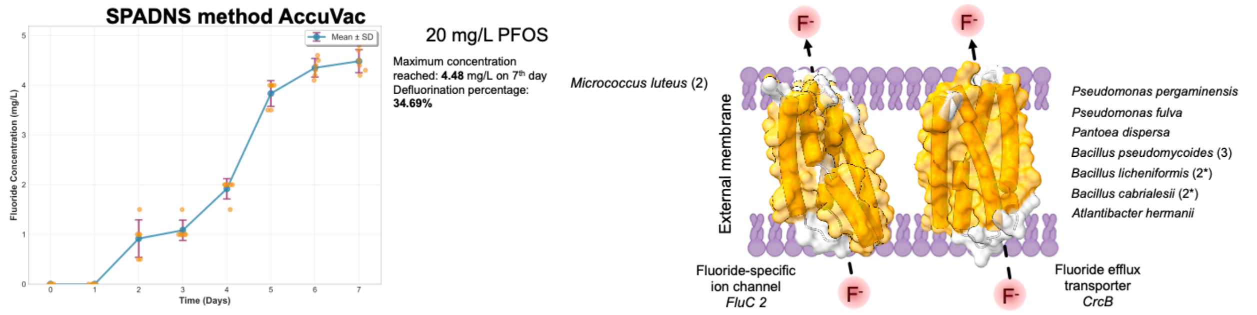


**Figure S2. Fluoride liberation kinetics and membrane transport mechanisms during PFOS defluorination by the GENIA consortium.** The left panel shows temporal fluoride release measured by the SPADNS method AccuVac, demonstrating biphasic kinetics with lag phase through day 1 followed by linear release (rate constant: 4.257 ± 0.333 mg $L^{-1}$ $d^{-1}$, $R^2 = 0.976$) reaching a maximum concentration of 4.48 mg $L^{-1}$ by day 7, corresponding to 34.69% defluorination efficiency. The right panel illustrates proposed membrane transport mechanisms for fluoride detoxification, featuring fluoride efflux transporter CrcB (present in 8/9 consortium strains, 88.9% coverage) and fluoride-specific ion channel FluC 2, enabling efficient $F^-$ export to prevent cytotoxic accumulation. The consortium member distribution shows broad representation across Pseudomonas, Bacillus, and other genera, with numbers in parentheses indicating gene copy numbers.
